# The Genetic Basis of Microbiome Recruitment in Grapevine and its Association with Fermentative and Pathogenic Taxa

**DOI:** 10.1101/2025.03.31.646331

**Authors:** Lena Flörl, Jerry Lin, Reid G. Griggs, Mélanie Massonnet, Noé Cochetel, Rosa Figueroa-Balderas, Dario Cantu, Nicholas A. Bokulich

**Affiliations:** Department of Health Sciences and Technology, ETH Zurich, Switzerland; Department of Viticulture and Enology, University of California Davis, Davis, California, USA; Stony Hill Vineyard, St. Helena, California, USA; Genome Center, University of California Davis, Davis, California, USA

**Keywords:** Microbiome, Plant-Microbiome-Interactions, QTL Mapping, Plant-Fungal-Bacterial Interactions, Mycobiome, Terroir

## Abstract

- While grapevine is an exceptional perennial model for studying host-microbiome interactions, the host genome’s role in microbiome assembly is often masked by environmental factors. This research provides a first insight into the genetic mechanisms shaping berry-associated microbial communities.
- Using QTL mapping in a newly established population of 140 F1-progeny grapevine genotypes in a complete random block design, we were able to control abiotic effects and investigate how the host genome influences grape berry-associated bacterial and fungal communities.
- We identify significant associations between various microorganisms and the grape genome, including pathogenic fungi such as *Botrytis* spp. and fermentative yeasts such as *Saccharomyces cerevisiae*. Many of these taxa map to the same genetic loci associated with plant immune responses, suggesting that specific genetic loci broadly influence microbial community assembly in fruits.
- Our findings demonstrate that grapevine genetics significantly shape the microbiome, even under varying environmental conditions; moreover, that broad, rather than known symbiont-specific mechanisms control microbial colonization of fruit, revealing an emergent “domino” effect with implications for plant-fungal-bacterial interactions. We provide a framework for understanding genotype-microbiome interactions in perennial plants, enabling future targeted experiments to establish causal relationships in microbiome recruitment and offer a potential avenue for breeding programs advancing sustainable viticulture.

## Introduction

Grapevine-associated microbial communities are shaped by a complex interplay of biotic and abiotic factors (Griggs *et al*., 2021). Emerging evidence indicates that site-specific environmental effects largely shape grapevine microbiome composition, with the host genotype playing a secondary role (Bokulich *et al*., 2014; Portillo *et al*., 2016). It is therefore challenging to disentangle the impact of the variety from environmental influences on assembled microbial communities. However, cultivars were already shown to harbor distinct microbial communities when comparing them within individual growing regions (Bokulich *et al*., 2014, 2016; Portillo *et al*., 2016; Marzano *et al*., 2016) or the same vineyard (Awad *et al*., 2023).

These observed differences in microbial communities associated with grape berries of different varieties may be influenced by variations in chemical composition, such as available nutrients, pH, and secondary metabolites. For example, red varieties contain higher levels of polyphenols, including anthocyanins, which have antimicrobial properties (Coman *et al*., 2018). Additionally, morphological traits like cluster density, skin thickness, and ripening patterns create distinct surface conditions that may influence microbial colonization. For instance, thin-skinned and tightly clustered varieties such as Pinot Noir and Zinfandel are more susceptible to sour rot of ripe berries (Brischetto *et al*., 2024). Similarly, the susceptibility of certain varieties to pathogens like *Botrytis* spp. has been associated with berry morphology (Herzog *et al*., 2015). Ultimately, these cultivar differences not only impact vine health but may also contribute to the uniqueness of the resulting wines via the microbiome, as native microbial communities produce metabolites that shape the chemical profile and flavor of the wine (Garofalo *et al*., 2015; Pinto *et al*., 2015).

However, the relationships between the host genome and associated microbial communities, as well as the specific genetic mechanisms underlying the assembly of these distinct communities, have not been explored yet. Here, we address for the first time open questions regarding how grapevine host genotype influences the berry microbiome. In a preliminary study, grape-associated microbial communities were characterized in several grape cultivars grown in mono-variety, adjacent blocks in a single vineyard with identical management practices. Samples were collected throughout each block, to evaluate inter-block variation and each variety was sampled before harvest as well as at optimal ripeness (Figure 1. A). In a subsequent study we collected berry samples from a newly established F1 population (Riesling x Cabernet Sauvignon) in a random block design for two vintages. All 140 different genotypes of this experimental population were genotyped, which allowed for associated microbial communities to be linked to specific genetic regions with quantitative trait loci (QTL) mapping (Figure 1. B). Since the sampled F1 genotypes differ in berry color, ripening behavior, and metabolomics, we anticipated differences in their microbial communities. Together, these studies examine intra-vineyard spatial and genotype effects. We show that microbial communities differ by location and cultivar, and show the first evidence of genotype-driven selection of grapevine microbiota, which could influence grapevine health and wine fermentation outcomes.

**Figure 1:**
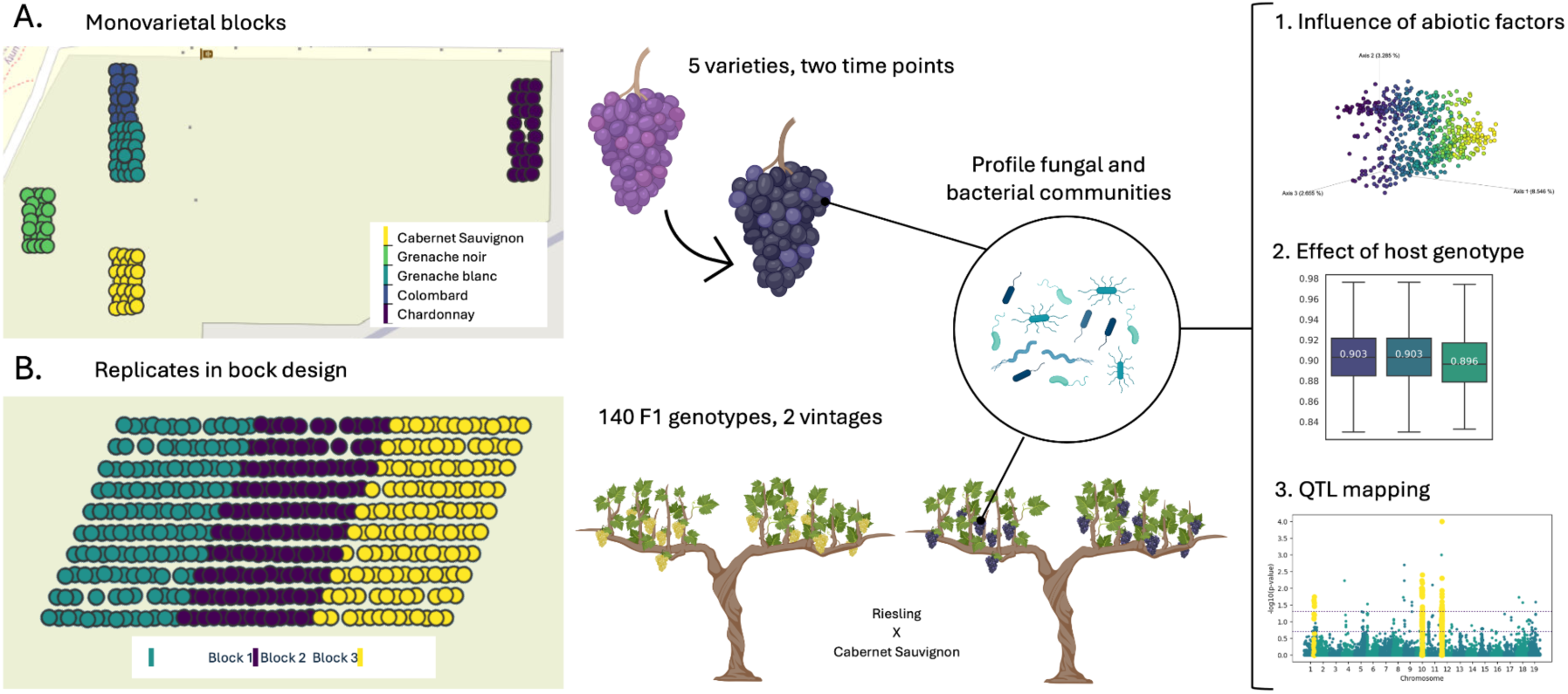
Experimental design of (A) the preliminary study collected in a UC Davis teaching vineyard and (B) the RxCS F1 population at the Oakville experimental station. In both studies, we profiled bacterial and fungal communities associated with grape berries. For the preliminary study berries were collected at two time points (pre-ripeness and peak ripeness) from multiple vines of five varieties within monovarietal blocks from a single vineyard. In the subsequent study, we sampled berries of two vintages from 140 F1 progeny of Riesling x Cabernet Sauvignon, which were planted in replicates across three randomized blocks.

## Results

### Different Varieties Harbor Distinct and Temporally Robust Microbial Communities

In the preliminary study, we aimed to test whether the geographic distance as well as grapevine variety appear to collectively structure microbial communities robustly throughout berry maturation. We characterized bacterial and fungal communities of grapes collected at commercial ripeness from 5 varieties planted in consecutive blocks within a single vineyard (Davis, CA, USA) using 16S rRNA gene and internal transcribed spacer (ITS) amplicon sequencing. This yielded a total of 182 samples and 3’888 unique amplicon sequencing variants (ASVs) with a total frequency of 241’360 for 16S rRNA gene amplification; and 122 samples, 6’168 ASVs and a total frequency of 2’557’136 for ITS. Permutational multivariate analysis of variance (PERMANOVA) tests of beta diversity metrics revealed that bacterial and fungal community structure was significantly different between varieties (Table 1 and Figure 2). Taking the abundance of microbial ASVs into account, the grapevine variety is shown to explain up to 57.1% of variation in bacterial communities (p_Bray Curtis_ = 0.001) and 20.7% in fungal communities, respectively (p_Bray Curtis_ = 0.001). Notably, while different time points significantly influenced fungal community composition (R2_Bray Curtis_ = 0.043, p_Bray Curtis_ = 0.001; R2_Jaccard_ = 0.013, p_Jaccard_ = 0.005), this effect becomes insignificant when accounting for variety (R2_Bray Curtis_ = 0.005, p_Bray Curtis_ = 0.384; R2_Jaccard_ = 0.009, p_Jaccard_ = 0.391). This indicates that, during the final weeks of ripening, variety exerts a stronger influence on microbial communities. Alpha diversity metrics also revealed significant differences in fungal community composition between the varieties (Kruskal-Wallis: p_Evenness_ < 0.001, p_Observed Features_ < 0.001, p_Shannon_ < 0.001). The lack of significant changes in fungal alpha diversity between time points (Kruskal-Wallis: p_Evenness_ = 0.104, p_Observed Features_ = 0.474, p_Shannon_ = 0.724) may be attributed to the relatively narrow sampling window, which spanned a maximum of 20 days. Similarly, bacterial alpha diversities differed significantly between varieties (Kruskal-Wallis: p_Evenness_ < 0.001, p_Observed Features_ <0 .001, p_Shannon_ < 0.001), however was also significantly different between the sampling time points (Kruskal-Wallis: p_Evenness_ = 0.0140, p_Observed Features_ < 0.001, p_Shannon_ = 0.001).

**Figure 2:**
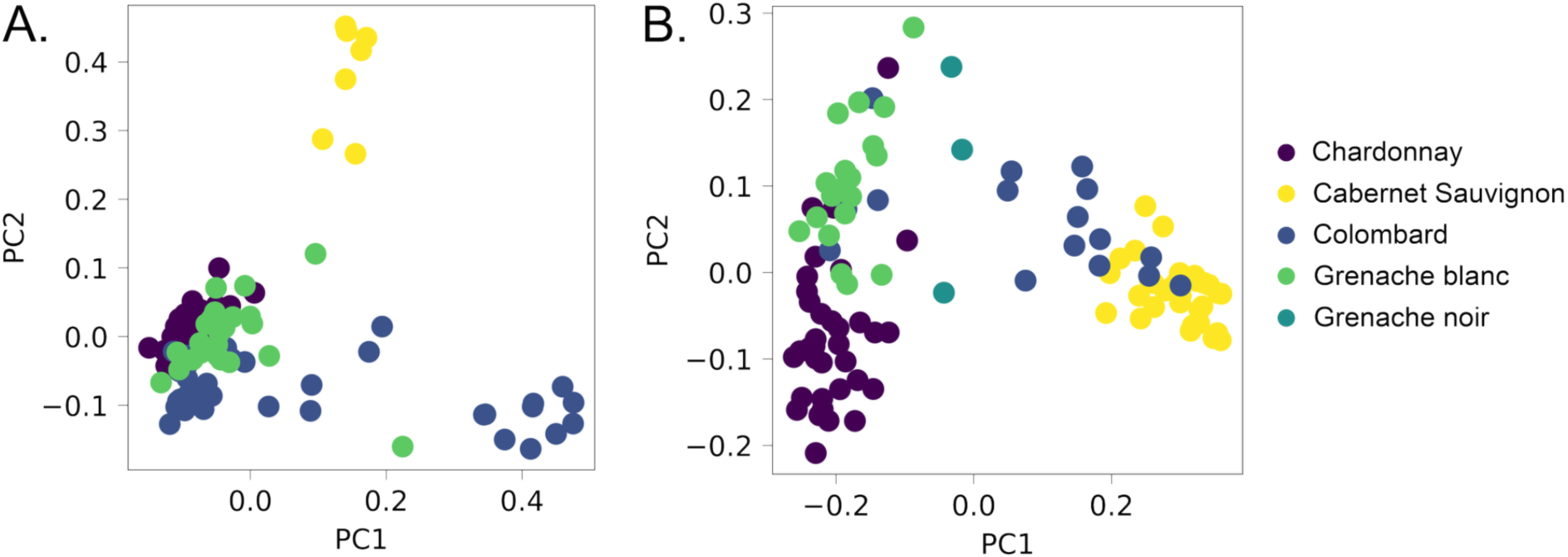
PCoA plots of the Jaccard beta diversity metrics for fungi (A), as well as for bacteria (B) showing that samples cluster by variety.

**Table 1:** Permutational multivariate analysis of variance (PERMANOVA) tests of bacterial and fungal diversity metrics with various covariates and their interactions reveal the significant effect of grapevine varieties in the monovarietal block experiment

|  | Bacteria |  |  |  | Fungi |  |  |  |
| --- | --- | --- | --- | --- | --- | --- | --- | --- |
|  | Bray Curtis |  | Jaccard |  | Bray Curtis |  | Jaccard |  |
|  | R2 | p-value* | R2 | p-value* | R2 | p-value* | R2 | p-value* |
| Variety | 0.571 | **0.001** | 0.175 | **0.001** | 0.207 | **0.001** | 0.088 | **0.001** |
| Vine | 0.045 | 0.107 | 0.079 | 0.084 | 0.079 | **0.006** | 0.082 | 0.099 |
| Vineyard Row | 0.012 | 0.297 | 0.025 | 0.370 | 0.042 | **0.001** | 0.032 | **0.005** |
| Time Point | 0.007 | 0.112 | 0.009 | 0.143 | 0.043 | **0.001** | 0.013 | **0.005** |
| Variety : Vine | 0.082 | 0.058 | 0.144 | 0.138 | 0.124 | **0.002** | 0.130 | **0.016** |
| Variety : Vineyard Row | 0.040 | 0.082 | 0.070 | 0.104 | 0.046 | **0.014** | 0.048 | **0.042** |
| Variety : Time Point | 0.012 | 0.115 | 0.017 | 0.214 | 0.005 | 0.384 | 0.009 | 0.391 |
| Vine : Time Point | 0.029 | **0.022** | 0.037 | 0.058 | 0.028 | 0.347 | 0.042 | 0.682 |
| Vineyard Row : Time Point | 0.024 | **0.009** | 0.027 | 0.117 | 0.115 | **0.001** | 0.042 | **0.001** |

\* p-values < 0.05 are considered significant and highlighted in bold

To evaluate spatial heterogeneity, we included the individual vine and the vineyard row from which the sample was collected in the PERMANOVA analysis of beta diversity metrics (Table 1). While bacteria show little spatial heterogeneity (Vine: R2_Bray Curtis_ = 0.045, p_Bray Curtis_ = 0.107; R2_Jaccard_ = 0.079, p_Jaccard_ = 0.084; Vineyard Row: R2_Bray Curtis_ = 0.012, p_Bray Curtis_ = 0.297; R2_Jaccard_ = 0.025, p_Jaccard_ = 0.370), fungal communities are significantly influenced by the location of sampling (Vine: R2_Bray Curtis_ = 0.079, p_Bray Curtis_ = 0.006; R2_Jaccard_ = 0.082, p_Jaccard_ = 0.099; Vineyard Row: R2_Bray Curtis_ = 0.042, p_Bray Curtis_ = 0.001; R2_Jaccard_ = 0.032, p_Jaccard_ = 0.005). Evidence of spatial relationships between fungal communities appears to be strengthened by significant Mantel tests between beta diversity metrics and geodesic distances when testing across all varieties (p_Bray Curtis_ < 0.001, p_Jaccard_ < 0.001). However, the monovarietal blocks are planted in separate locations throughout the vineyard (see Figure 1), with Chardonnay samples specifically positioned at the far northern end. Thus, performing Mantel tests within the individual varieties revealed no significant intra-block variation, except for Chardonnay samples (Fungi: Grenache blanc: p_Bray Curtis_ = 0.133, p_Jaccard_ = 0.113; Cabernet Sauvignon: p_Bray Curtis_ = 0.841, p_Jaccard_ = 0.693; Colombard: p_Bray Curtis_ = 0.339, p_Jaccard_ = 0.643; Chardonnay: p_Bray Curtis_ = 0.001, p_Jaccard_ = 0.936).

To further disentangle the effect of varieties and spatial patterns, we controlled for phylogenetic relatedness of sampled genotypes while testing for inter-varietal effects using partial Mantel tests. This is particularly interesting as some varieties, Cabernet Sauvignon with Colombard, and Grenache Blanc with Grenache Noir, are clonal to each other (Supplementary Figure 1). The partial Mantel tests controlling for phylogenetic distance (microbial distance matrix ∼ geodesic distance + phylogenetic distance) shows that microbial diversity is still significantly shaped by spatial effects within the vineyard (Bacteria: Mantel R_Bray Curtis_ = 0.1222, p_Bray Curtis_ = 0.003, R_Jaccard_ = 0.1574, p_Jaccard_ = 0.001; Fungi: Mantel R_Bray Curtis_ = 0.1217, p_Bray Curtis_ = 0.002, R_Jaccard_ = 0.0855, p_Jaccard_ = 0.001). Inversely, when controlling for spatial effects, the partial Mantel test also reveals that phylogenetic relatedness between grapevine varieties (microbial distance matrix ∼ phylogenetic distance + geodesic distance) significantly influences both the abundance and composition of bacterial communities, as well as the composition of fungal communities (Bacteria: Mantel R_Bray Curtis_ = 0.1861, p_Bray Curtis_ = 0.006, R_Jaccard_ = 0.1881, p_Jaccard_ = 0.005; Fungi: Mantel R_Bray Curtis_ = 0.193, p_Bray Curtis_ = 0.193, R_Jaccard_ = 0.1414, p_Jaccard_ = 0.0036). This suggests that the differences observed between varieties in bacterial and fungal community compositions is driven by cultivar-specific (i.e., genotype) effects, as well as by spatial (e.g., block) effects; this is a sensible outcome, given the small size and homogeneity (management, soil profile, topography) of the vineyard site tested here.

This preliminary study robustly showed that varieties harbor distinct microbial communities, even at a close range, in mono-varietal blocks within a single vineyard. However, the dataset has two main limitations: the lack of sampling on the same day, as sampling was conducted based on ripeness stage, and the lack of spatial replication. Finally, this experiment demonstrates only that the grapevine microbiota differs in commercial varieties, but does not provide insight into the genetic factors underpinning this relationship. To address these limitations and further investigate the specific effects of host genotype on microbiome assembly, we conducted a subsequent study in an experimental F1 population.

### Microbial Communities on Berries of the Same Genotype Are More Similar to Each Other Even Than Between Closely Related Genotypes

Next, we sought to more closely investigate how grapevine genotype influences the grape microbiome in an experimental vineyard setting. For two years we characterized the grape berry microbiome of a newly established experimental population of 1’260 vines, with 140 different F1 genotypes deriving from the cross of Riesling and Cabernet Sauvignon (hereafter RxCS), each planted in triplicate across three blocks in complete random design (see Figure 1.B). To disentangle the effects of different sampling strategies, berries were collected at the same ripeness level (24 °Brix) in 2020, and to parse climatic effects, sampling was conducted at the same time point in 2022. Given the high climate sensitivity of grapevines, direct comparison of microbial communities across vintages is inherently challenging. As detailed in Supplementary Table 1, the daily maximal and minimal temperatures varied as expected substantially between the sampling periods in each year. The 2020 season showed a higher cumulative heat exposure, reaching a total of 3’556.75 °F growing degree days (GDD), compared to 3’363.2 °F GDD in 2022, underscoring the distinct environmental conditions each year.

We performed ultra-deep 16S rRNA gene and ITS domain amplicon sequencing on crushed grape samples, yielding 17’166 unique bacterial ASVs and a total count of 12’696’835 sequences and 46’777 fungal ASVs and a total count of 203’982’580 sequences. To reveal factors that influence bacterial and fungal community structure in this experimental vineyard we first studied differences in alpha diversity. The block from which samples were collected did not have a significant effect on most alpha diversity metrics, except for evenness and Shannon entropy for bacteria (Kruskal-Wallis for Bacteria: p_Evenness_ = 0.018, p_Observed Features_ = 0.080, p_Shannon_ = 0.0247, p_Faith’s Phylogenetic Diversity_ = 0.073; Fungi: p_Evenness_ = 0.071, p_Observed Features_ = 0.270, p_Shannon_ = 0.699). Both, the bacterial and fungal samples from 2020 had a significantly higher alpha diversity than 2022 (Kruskal-Wallis for Bacteria: p_Evenness_ = 6.90e-12, p_Observed Features_ = 9.51e-29, p_Shannon_ = 2.65e-9, p_Faith’s Phylogenetic Diversity_ = 2.44e-30; Fungi: p_Evenness_ = 2.66e-15, p_Observed Features_ = 1.45e-32, p_Shannon_ = 3.45e-9), which is expected as communities are known to increase in population size over the ripening process (Griggs *et al*., 2021) and the 2020 samples were all collected at peak ripeness whereas there was more heterogeneity in 2022. To take these differences in sampling strategies and their impact on differences between years into account, we included the Shannon alpha diversity index in the subsequent PERMANOVA of beta diversity metrics (Table 2).

**Table 2.**
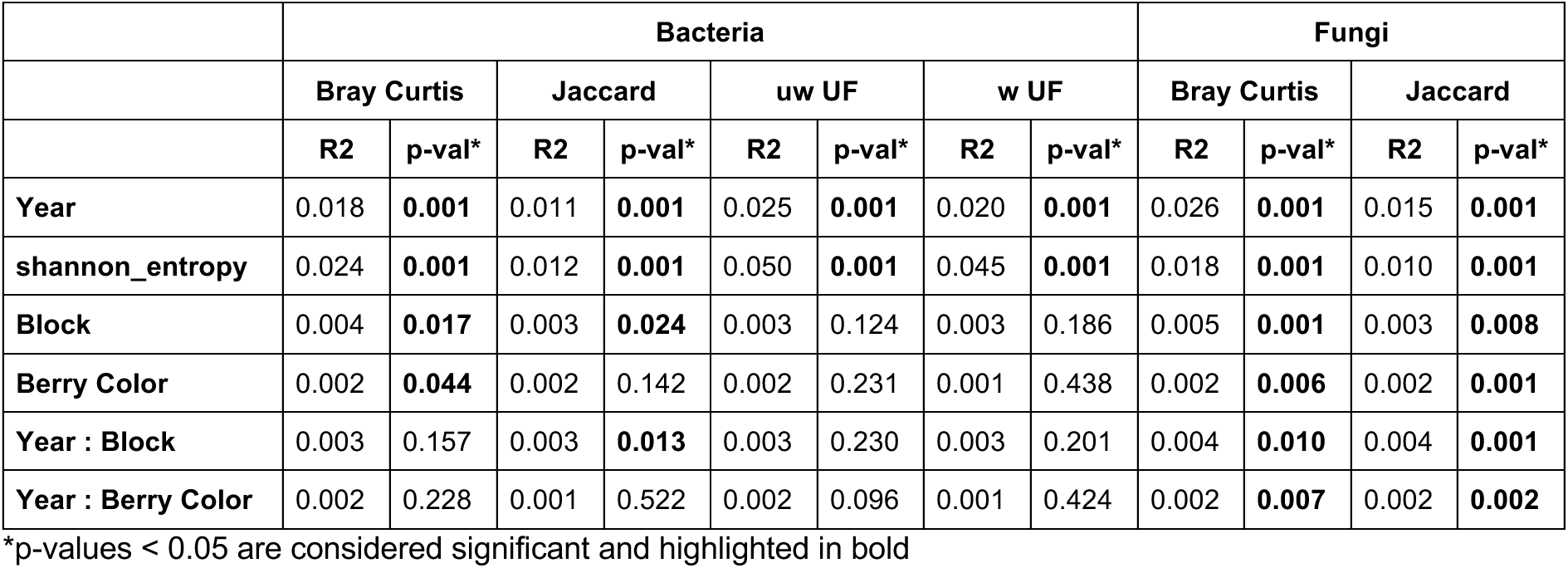
PERMANOVA analyses of fungal and bacterial data in the F1-cross vineyard reveal that beta diversity is primarily driven by differences between years and is also influenced by alpha diversity (Shannon entropy).

|  | Bacteria |  |  |  |  |  |  |  | Fungi |  |  |  |
| --- | --- | --- | --- | --- | --- | --- | --- | --- | --- | --- | --- | --- |
|  | Bray Curtis |  | Jaccard |  | uw UF |  | w UF |  | Bray Curtis |  | Jaccard |  |
|  | R2 | p-val* | R2 | p-val* | R2 | p-val* | R2 | p-val* | R2 | p-val* | R2 | p-val* |
| **Year** | 0.018 | **0.001** | 0.011 | **0.001** | 0.025 | **0.001** | 0.020 | **0.001** | 0.026 | **0.001** | 0.015 | **0.001** |
| **shannon_entropy** | 0.024 | **0.001** | 0.012 | **0.001** | 0.050 | **0.001** | 0.045 | **0.001** | 0.018 | **0.001** | 0.010 | **0.001** |
| **Block** | 0.004 | **0.017** | 0.003 | **0.024** | 0.003 | 0.124 | 0.003 | 0.186 | 0.005 | **0.001** | 0.003 | **0.008** |
| **Berry Color** | 0.002 | **0.044** | 0.002 | 0.142 | 0.002 | 0.231 | 0.001 | 0.438 | 0.002 | **0.006** | 0.002 | **0.001** |
| **Year : Block** | 0.003 | 0.157 | 0.003 | **0.013** | 0.003 | 0.230 | 0.003 | 0.201 | 0.004 | **0.010** | 0.004 | **0.001** |
| **Year : Berry Color** | 0.002 | 0.228 | 0.001 | 0.522 | 0.002 | 0.096 | 0.001 | 0.424 | 0.002 | **0.007** | 0.002 | **0.002** |

\*p-values < 0.05 are considered significant and highlighted in bold

As anticipated, the bacterial and fungal communities varied significantly between the years, yet this only accounted for a relatively small portion of the variance explained (1.1 - 2.6%). The block had a similarly small effect, explaining 0.3 - 0.5% of the variance, which was significant for fungal communities but only for bacterial communities when disregarding phylogenetic distances between taxa. Notably, the effect of the block remained significant for fungal communities even when nesting the effect within the year (Table 2). However, subsequent Mantel tests correlating microbial composition differences with spatial distances between individual grapevines did not reveal any substantial spatial heterogeneity, except for some minor differences in microbial abundance in 2022 (Mantel Fungi: p_Bray Curtis, 2020_ = 0.233, p_Jaccard, 2020_ = 0.308, p_Bray Curtis, 2022_ = 0.006, p_Jaccard, 2022_ = 0.761; Bacteria: p_Bray Curtis, 2020_ = 0.747, p_Jaccard, 2020_ = 0.589, p_weighted UniFrac, 2020_ = 0.612, p_unweighted UniFrac, 2020_ = 0.533, p_Bray Curtis, 2022_ = 0.698, p_Jaccard, 2022_ = 0.977, p_weighted UniFrac, 2022_ = 0.048, p_unweighted UniFrac, 2022_ = 0.705).

Interestingly, the berry color appeared to modestly, yet robustly impact assembled fungal communities, while it was only significantly associated with differences in bacterial communities for the Bray Curtis metric (Table 2).

Having demonstrated significant differences by sampling year in microbial communities with minimal spatial variation across the blocks, we then investigated the influence of host genotype on associated microbiomes. A comparison of intra- and inter-genotype beta diversity metrics confirmed that communities on vines of the same genotype were significantly more similar to each other (see Figure 3). This was robust across both years for all beta diversity metrics of bacterial and fungal communities (see Supplementary Table 2). This is the first clear demonstration that, when controlling for spatial and temporal effects, even closely related grapevine varieties select for distinct microbial communities.

**Figure 3.**
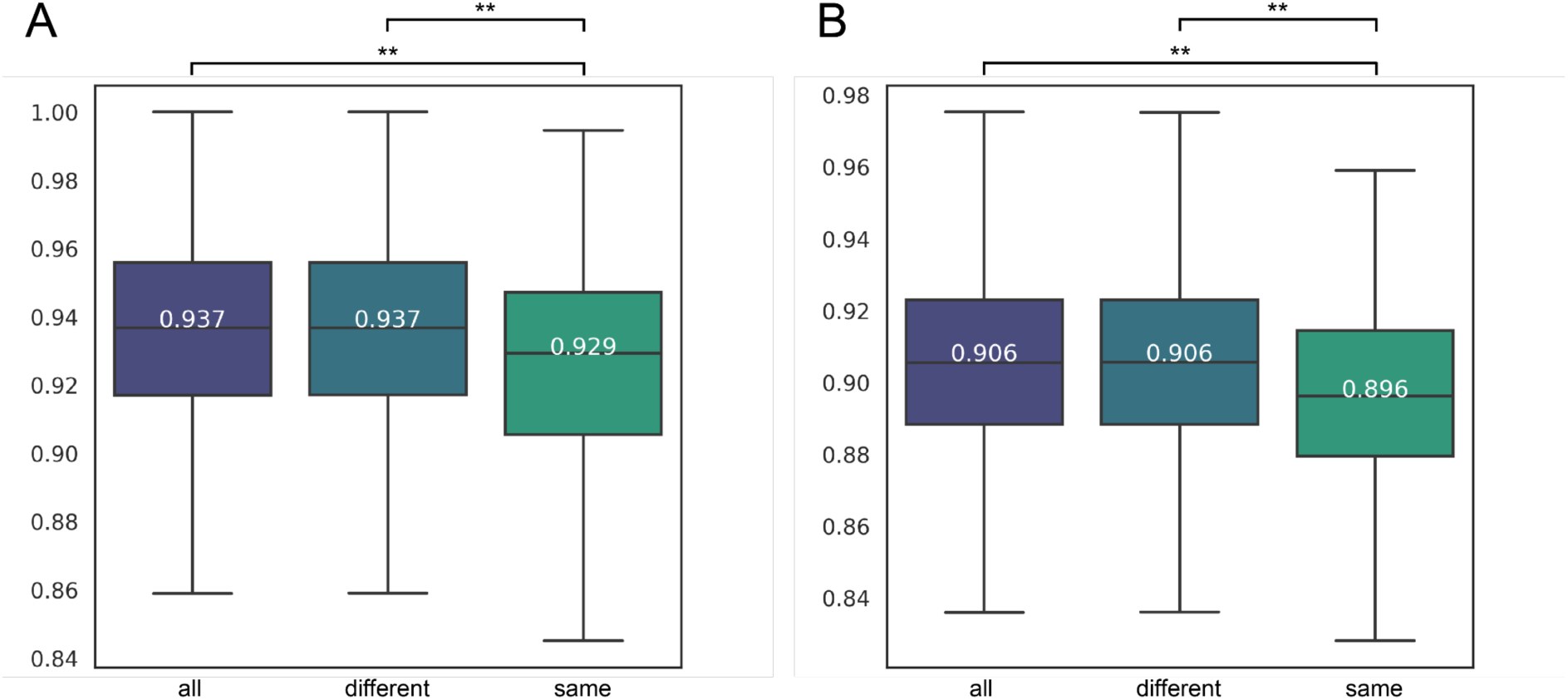
Comparing Jaccard distances of 2020 samples for bacteria (A) and fungi (B). The corresponding Kruskal-Wallis tests are calculated for the distances of all samples (purple), samples of different genotypes (blue) and samples of the same genotypes (green). The significantly lower Jaccard distances (** indicating a p value < 0.001) of samples with the same genotype, reveal that microbial communities are more similar on grapevines with the same genetic background.

### Microbial Features Are Significantly Associated With Distinct Genetic Loci

Building on our findings that grapevines of the same genotype harbor distinct microbial communities, we next investigated the genetic mechanisms driving this selective recruitment through QTL mapping. First, we built a genetic map using the haplotype 1 of the Cabernet Sauvignon genome and genotype-by-sequencing data from the 140 F1 genotypes of the RxCS population. In total we identified 5’386 markers across the 19 chromosomes with a density of 28.96 ± 22.25 cM and a genotype completion of 97.8% (Supplementary Figure 2). To study the microbial features as quantitative traits, the ASVs were clustered at 99% similarly into OTUs (Operational Taxonomic Units) and rare features were filtered out. Each dataset was subset by year and the median feature count per genotype from samples across all blocks were calculated. To study features at various taxonomic levels, OTUs were further collapsed to genus, family, and order level. Subsequently, these were converted to presence/absence matrices to reduce noise in the data. Ultimately we performed interval mapping with a Haley– Knott regression to retrieve microbe-associated QTLs from a total of 10’669 features, resulting in 202’711 QTLs. We considered QTLs to be significant only with a logarithm of the odds (LOD) score above the significance threshold (LOD > 4.9) as well as with a genome-wide adjusted p-value below 0.05. A significance threshold for the LOD was established with 1,000 permutations at a 95% confidence interval to ensure the null hypothesis of no QTL at a given marker holds true during interval mapping. This yielded a total of 192 significant QTLs (see Supplementary Table 3), the majority of which were linked to the same set of markers on chromosome 5 (see Supplementary Table 4), creating a distinct hotspot that is strongly associated with multiple taxa (see Figure 4).

**Figure 4.**
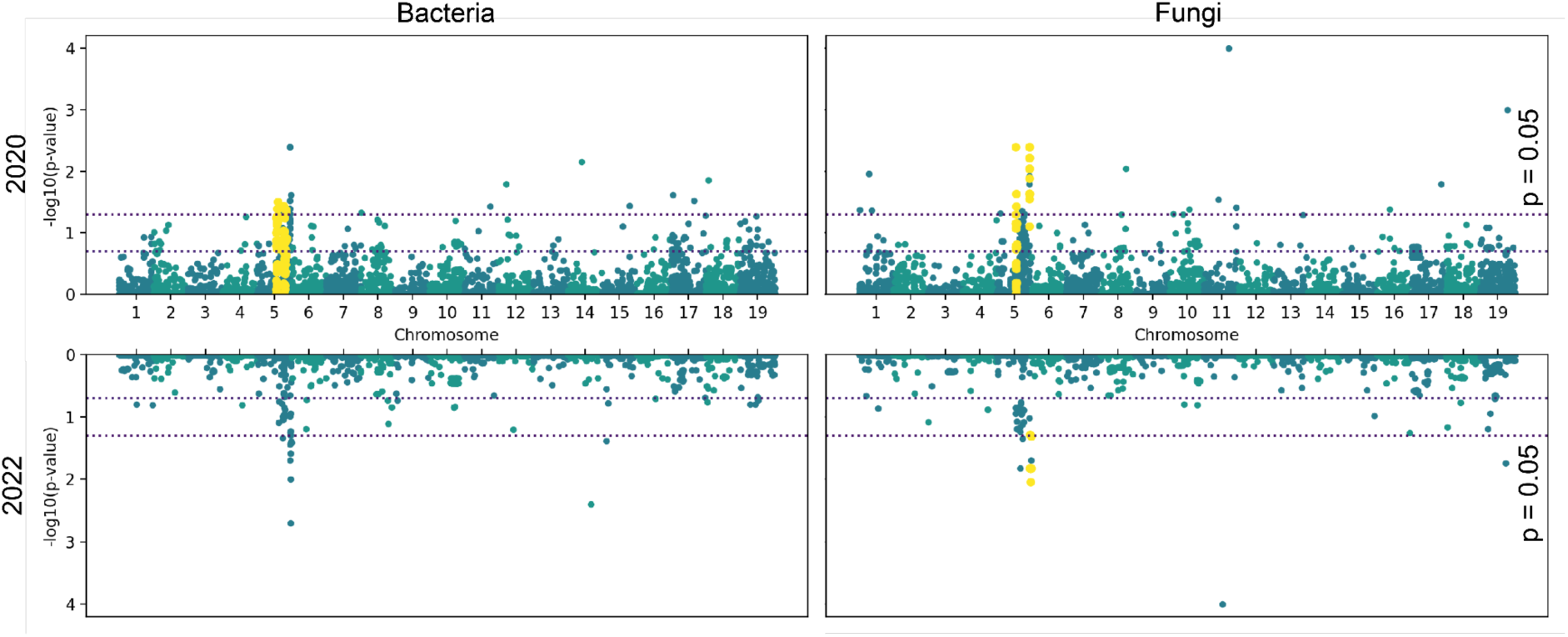
Manhattan plots of all QTLs across the 19 grape chromosomes and the corresponding -log10(p-value) from mapping bacterial (left) and fungal (right) OUTs for the years 2020 (top) and 2022 (bottom) reveal a hotspot on chromosome 5 associated with multiple microbial features (highlighted in yellow).

Several genera contain OTUs that consistently yield significant QTLs across both years, with the majority mapping to markers within the chromosome 5 hotspot. Among fungal OTUs, these include *Filobasidium* spp. (specifically *Filobasidium chernovii), Cladosporium* spp. (specifically *Cladosporium tenuissimum* and *Cladosporium velox), Epicoccum nigrum, Vishniacozyma* spp. (specifically *Vishniacozyma victoriae)* and with lower LOD scores also *Mycosphaerella tassiana* and *Alternaria* spp. These fungal genera have been described as members of the typical ‘core’ grape berry microbiome (Liu & Howell, 2021). Similarly, bacterial OTUs from the genera *Lysinibacillus* (formerly *Bacillus*) and *Bacillus* also exhibit significant QTLs in both years, primarily mapping to the chromosome 5 hotspot. Strains of both genera have been shown to possess antifungal properties and have been suggested as potential plant growth promoting agents (Andreolli *et al*., 2016; Pacifico *et al*., 2019; Veras *et al*., 2023), while some *Lysinibacillus* strains were also reported to have grape spoilage potential (Gao *et al*., 2020).

### Interval Mapping Reveals Significant QTLs Associated with Opportunistic and Pathogenic Fungi

Many of the significant QTLs for fungal OTUs are associated with putative plant pathogens (2022: 17 out of 27, 2022: 16 out of 47), which are also among the features with the highest LOD scores. Mapping to the same chromosome 5 hotspot marker CHR05_24281346 in 2020 with high LOD scores (between 11.2 and 250.6) and sharply defined support intervals (Bayes Credible Interval (BCI): 3.36e-04 to 4.00e-06 cM) are different OTUs of *Botrytis* spp., *M. tassiana,* and *E. nigrum*. Multiple OTUs of the latter, *M. tassiana* and *E. nigrum*, as well as *Neocatenulostroma microsporum* and the genus *Fusarium* are also mapping to a marker on chromosome 5 in 2022 (CHR05_24608659, LOD: 129.4, BCI: 77-81 cM). This robust and strong association with the host genotype of *M. tassiana* and *E. nigrum* in both years is interesting, as both are well described grapevine commensal endophytes and opportunistic pathogens as saprophytes during senescence, i.e. as decomposers for decaying plant matter (Grube *et al*., 2011; Liu *et al*., 2020). *M. tassiana* is also a common genera associated with grapevine trunk disease (Patanita et al. 2022) as well as a core species in non-symptomatic plants (Liu *et al*., 2020). Some strains of *E. nigrum*, a genotypically and phenotypically highly variable species, are applied as biological control agents in various crops due to its production of secondary metabolites with antifungal and antibacterial properties (Del Frari *et al*., 2019). Particularly noteworthy is the strong association in 2020 with *Botrytis* spp. (CHR05_24281346, LOD: 250.6, BCI: 4e-6 cM), a well known grapevine pathogen causing bunch rot in berries (Bhatia *et al*., 2020). *Botrytis* acts as a necrotroph (i.e. inducing host cell death) and has therefore commonly been described to co-occur with saprophytes (Rienth *et al*., 2021), as we also show here with *M. tassiana* and *E. nigrum* which map to the same marker.

Other fungal pathogens yielding significant QTLs in 2020 include the obligate pathogens *Melampsora* spp. (CHR19_17331846, LOD: 6.2, BCI: 1.2 cM) and *Ustilago* spp. (CHR17_11666180, LOD: 5.2, BCI: 21.1 cM). OTUs of both also form central nodes in the respective network of fungi (see Section below and Figure 5). While *Melampsora* spp. are reported to correlate with other grapevine leaf rust pathogens (Ono *et al*., 2012), *Ustilago* spp. as well as *Devriesia* spp., mapping in 2022 (CHR05_22428041 LOD: 11.6, BCI: 81 cM) have been reported to co-occur with other pathogens in grapevine trunk disease (Adejoro *et al*., 2023).

**Figure 5.**
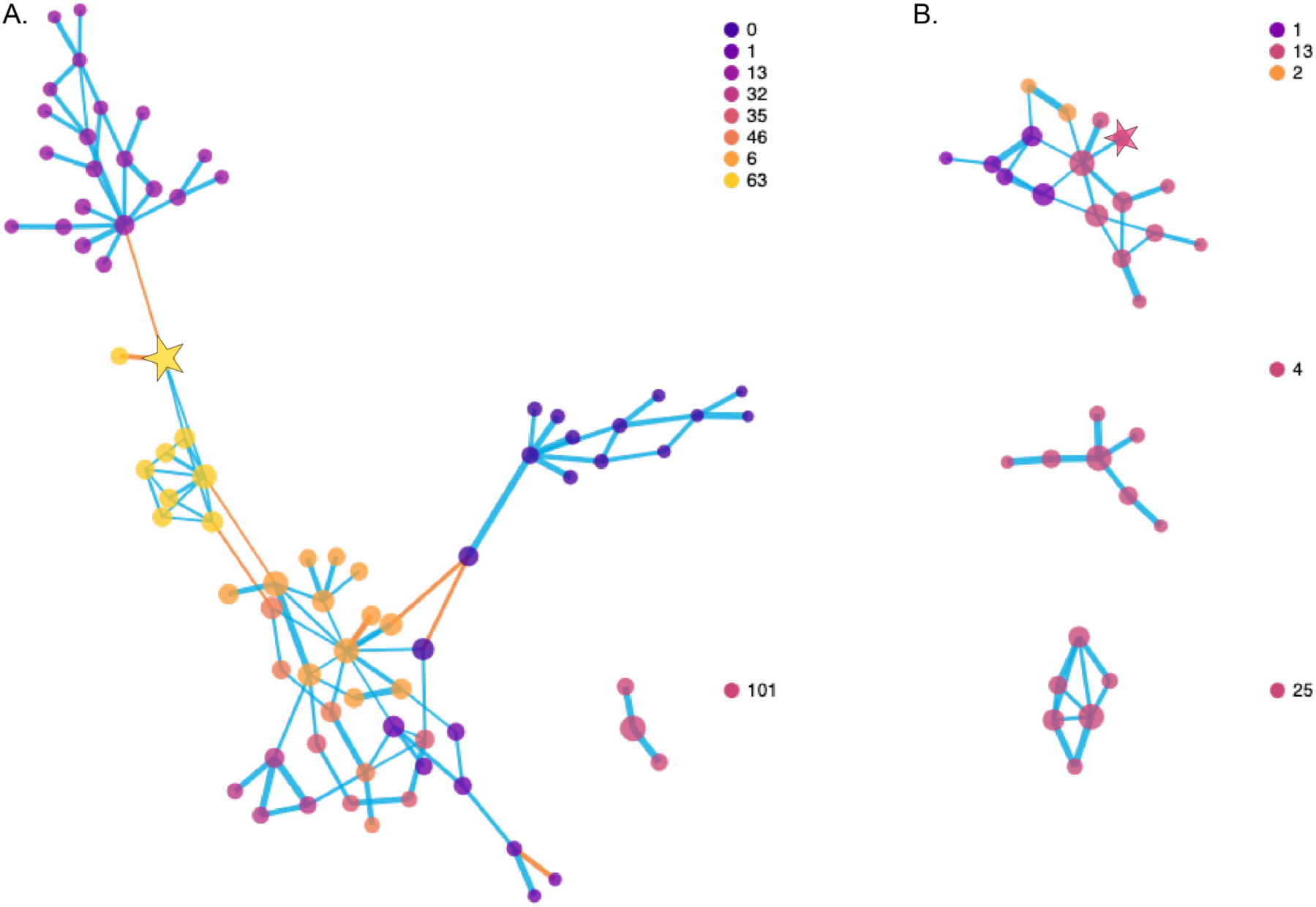
Visualization of the network analysis showing the relationships between fungal and bacterial features for 2020 (A) and 2022 (B). Each node is representing an OTU, with node size scaled by closeness of centrality and colored by community affiliation. The network edges are colored by correlation (blue for positive and orange for negative) and the thickness of edges is scaled according to the strength of connection between features. Network analysis of features in 2020 yielded 2 distinct groups and a total of 220 communities (including pairs and singletons), and 3 groups and 64 communities in 2022. The yeast *S. cerevisiae* is highlighted as a star in both years. These network visualizations but colored by taxonomic affiliation on the order level are shown in Supplementary Figure 3.

Our findings show that putative fungal pathogens exhibit strong associations with the host genotype, with many mapping to the same genetic locus. Notably, some of these putative pathogens are opportunistic and common commensals in healthy vines. Their co-occurrence of commensals with obligate pathogens suggests that the contribution may contribute to disease development. Further, these associations appear to be robust, as they are consistent across both years, despite differences in sampling strategies.

### The Genotype Impacts Fermentative Yeast Presence and Suggests Potential Pathogen Interactions

Given the impact of grape microbiota on subsequent wine production, the presence of potentially fermentative yeasts on the grape berry surface is of high interest. In 2020, the berries were collected at the same ripeness stage, allowing for the genotype to exert selective pressure in recruiting specific fermentative microbes. Mapping to the same chromosome 5 hotspot was an OTU of the key fermentative yeast *Saccharomyces cerevisiae* (CHR05_25035791, LOD: 9.9, BIC: 0.6 cM) as well as for *Rhodotorula babjevae* (CHR05_24608659, LOD: 19.5, BCI 3.4e-4 cM). Different fermentative organisms, including distinct strains of *S. cerevisiae*, have been previously isolated from various grapevine varieties (Blanco *et al*., 2006; Cordero-Bueso *et al*., 2011). However, to our knowledge, this is the first study to demonstrate the direct role of the host genotype in recruiting specific fermentative organisms.

Further, it is noteworthy that both yeasts are associated with a hotspot which is also linked to putative pathogens, including *Botrytis* spp. This association may arise from pathogen-induced damage to the berry surface, which releases sugars that support yeast growth. While *Saccharomyces* spp. were detected in the absence of *Botrytis*, all genotypes in 2020 where *Botrytis* were detected also contained *Saccharomyces* spp. Viable strains of *S. cerevisiae* can persist in *Botrytis*-infected berries, where they interact with *Botrytis cinerea* through both antagonistic mechanisms, such as growth and contact inhibition, or nutrient competition, and synergistic interactions like cross-feeding (Di Canito *et al*., 2021). Some strains of *S. cerevisiae* have thereby shown significant antagonistic effects and are being tested as biocontrol agents against phytopathogenic fungi like *B. cinerea* (Nally *et al*., 2012; Di Canito *et al*., 2021). Notably, the genus *Rhodosporidiobolus*, which maps to the same locus as *S. cerevisiae* and other pathogens in 2020, includes strains that, like *Saccharomyces*, can inhibit the growth of pathogens such as *Aspergillus* and *Penicillium* in grapes through the same mechanism of lytic enzyme production (Di Canito *et al*., 2021).

Here, we showcase the role of the host genotype in recruiting distinct fermentative yeasts, including *S. cerevisiae* and *R. babjevae*. These yeasts map to a genetic hotspot also associated with putative pathogens, suggesting a complex interaction between yeast recruitment, pathogen dynamics, and potential biocontrol mechanisms in grapevine health and wine production.

### Locus analysis reveals genes associated with plant host defense

The interval mapping reveals a clear hotspot of bacterial and fungal OTUs on chromosome 5, spanning an estimated 2.6 Mbp interval and containing 229 predicted genes. While GO terms enrichment analysis did not identify any enriched functions within this interval, the robustness of associations across two years, combined with high LOD scores and sharply defined support intervals, suggests that these may represent distinct loci directly influencing individual taxa.

Of particular interest is the marker CHR05_24281346, which yields significant QTLs for bacteria and fungi in both years and is strongly associated with numerous putative pathogens in 2020, including *M. tassiana*, *E. nigrum*, and *Botrytis* spp. The narrowly defined support interval for these significant QLTs directly maps to a class IV chitinase precursor (g473920). The expression of chitinase plays a key role in the defense of grapevine against fungi, specifically against *Botrytis* spp. (Salzman *et al*., 1998; Trotel-Aziz *et al*., 2006). Similarly, CHR05_24608659, associated with multiple pathogens in 2022 and the fermentative yeast *Rhodotorula babjevae* in 2020, maps to a chloroplastic-like cysteine synthase (g473630). Cysteine synthase is upregulated in plants in response to stress and pathogens, producing cysteine for synthesizing defense metabolites like glutathione, camalexin, and glucosinolates (Romero *et al*., 2014). It is noteworthy that two of the most robust and strong associations map to chitinase-related genes, which play a key role in degrading chitin in fungal cell walls, thereby releasing nutrients for other microbes (Shobade *et al*., 2024). This may explain the connection to bacterial genera such as *Lysinibacillus* and *Bacillus*, both of which map to the chromosome 5 hotspot and which themselves produce antifungal compounds like chitinases (Ahsan & Shimizu, 2021). This finding suggests that alleles related to chitinase activity could enhance plant defense against fungal pathogens, as seen in other plants, such as maize, where chitinase QTLs are considered promising for breeding against *Aspergillus* (Hawkins *et al*., 2015).

Another notable marker, CHR05_25035791, associated with fungal pathogens like *Alternaria* spp. in 2022 and the yeast *S. cerevisiae*, maps to an ankyrin-repeat-containing gene (g569720). Ankyrin repeats mediate protein–protein interactions that trigger pathogen recognition and defense signaling, activating plant immune responses (Kolodziej *et al*., 2021). These associations suggest that this gene may enhance microbial recognition or signal transduction, potentially contributing to increased resistance against pathogens. Overexpression of ankyrin-repeat genes has been shown to increase resistance to fungi in other plant hosts (Mou *et al*., 2013; Ngaki *et al*., 2016), making them potential breeding targets for pathogen resistance (Wang *et al*., 2020).

Overall the analysis of predicted genes located within the QTL intervals defined by significant markers reveals several genes associated with host defense proteins, which offer potential targets for breeding increased disease resistance. These findings also provide insight into potential mechanisms that could regulate microbial community composition, including the co-occurrence of commensals, fermentative yeasts, and pathogenic fungi in grapes.

### Network Analysis of Bacterial and Fungal Communities Uncovers Robust Communities Across Different Years

To further investigate these co-occurrences, we performed a network analysis of bacterial and fungal OTUs for each year (see Figure 5). In 2020, a distinct bacterial community formed around OTUs from the genera *Lysinibacillus* and *Bacillus* (community 13), alongside other common soil bacteria such as *Thermoactinomyces*, *Geobacillus*, *Romboutsia*, and *Turicibacter*. This community was negatively correlated with *S. cerevisiae*, a key fermentative yeast (highlighted as a star in the graph). *S. cerevisiae*, which yields a significant QTL at the marker CHR05_25035791, is positively correlated with a separate community (community 63) containing bacteria from *Pseudomonas*, *Sphingomonas*, *Bacillus*, *Enterococcus*, *Listeria*, *Staphylococcus*, *Lactobacillus fermentum*, and *Enterobacteriaceae*. Some of these bacteria, such as *Lactobacillus fermentum*, have been previously reported to co-occur with *S. cerevisiae* (Carvalho *et al*., 2020), and certain *Pseudomonas* strains can even support its growth (Romano & Kolter, 2005). At the heart of the 2020 network are multiple closely connected fungal communities. Many OTUs of which appear to be influenced by the plant genotype, including *Alternaria* spp. (with a significant QTL at CHR08_18287104), *Bipolaris* spp. (CHR01_533750), *Phialocephala* spp. (CHR10_12072879), *Microstroma* spp. (CHR11_6045134), and *Ustilago* spp. (CHR17_11666180). This is particularly evident in the closely connected OTUs from *Cladosporium* (community 32), *Mycosphaerella* (communities 32 and 35) and *Epicoccum* (community 35) which map to the same genetic locus (CHR05_24281346). Interestingly, *Mortierella* spp. also associates with this locus, but is negatively correlated and branches off to form an entirely separate community (community 0), together with OTUs from genera like *Penicillium*, *Umbelopsis*, *Hyaloscypha*, *Geminibasidium*, *Sagenomella*, *Phialocephala*, and *Paratritirachium*.

In 2022, the network analysis of fungal and bacterial OTUs revealed three distinct groups, some of which resembled the 2020 network. In community 13, *S. cerevisiae* was positively correlated with endophytes from the taxa *Mycosphaerella spp.*, *Stemphylium spp*., and *Microstroma* spp. Community 25, which resembled community 63 from 2020, contained *Lactobacillus fermentum*, *Enterococcus spp.*, *Listeria spp.*, *Staphylococcus spp.*, and *Bacillus* spp. Meanwhile, community 4, which resembled community 13 from 2020, included the typical soil bacteria like *Bacillus* spp., *Turicibacter* spp., and *Romboutsia* spp.

Our analysis reveals that certain fungal and bacterial communities are consistently present across the years, despite differences in ripeness and climatic factors, and are strongly associated with specific QTLs of the grapevine host. This suggests a stable host-microbe interaction, potentially driven by the grapevine’s genetic influence on the recruitment and maintenance of these microbial communities. Moreover, this suggests that host-microbiome interactions have additional “domino effects” whereby suppression of one clade may lead to emergence of a separate clade (e.g., presence of anti-fungal genes leads to increased growth of bacterial clades in the absence of fungal competitors or vice versa).

### Microbial Diversity Metrics are Modestly Associated With The Genotype

In addition to the recruitment of individual microbial taxa and given the strong community formation, we investigated the influence of the host genotype on microbial diversity. Therefore, interval mapping was performed using alpha diversity indices as well as the first two components of a principal coordinate analysis (PCoA) of beta diversity as quantitative traits. This yielded a significant QTL for the second principal component of the weighted UniFrac metric for bacterial communities in 2022 on chromosome 1 (CHR01_7538491, LOD: 4.9, p-val: 0.020) (see Supplementary Figure 4 and Supplementary Table 4). The weighted UniFrac metric, which takes phylogenetic distance between taxa and their abundance into account, varied considerably between the different alleles, and the QTL explained 18.5% of the variation (see Supplementary Figure 5). Gene Ontology (GO) enrichment analysis of the genes retrieved from the large support interval (16.9 cM) showed a small, yet significant positive enrichment for cellular homeostasis functions (see Supplementary Table 4). While the first component of the weighted UniFrac PCoA for bacteria was found to be significantly influenced by the differences in Shannon entropy between samples (see Table 2), the second component was mostly associated with the presence of *Bacillus* spp. OTUs, among others (see Supplementary Figure 6.B). Interestingly, the second component of the weighted UniFrac in 2022, which also yielded a suggestive QTL (LOD: 4.17, p-val: 0.084), is similarly influenced by different *Bacillus* spp. OTUs (see Supplementary Figure 6.A). In both years, *Bacillus* spp. OTUs form connective hubs in the networks and yield multiple significant QTLs (see Supplementary Figure 3).

Additionally, we recovered seven suggestive QTLs (LOD > 3.5, p-val < 0.2) from mapping diversity metrics (see Supplementary Table 5). The GO enrichment analysis of the respective support intervals reveals enrichment or depletion of key functions that could regulate the interaction between host and microbiome, such as response to nucleotide and protein binding (which includes for example the previously described ankyrin-repeat proteins), abiotic stimulus, signaling receptor activities or secondary metabolite processes (see Supplementary Table 6).

Although the results from mapping diversity metrics should be interpreted with caution due to the inherent noise in the data, they provide valuable insights suggesting that certain genetic regions may be linked to overall microbiome assembly and potential functions underlying this association. Further, the results suggest that genetic associations appear to be influenced by microbes central to microbial networks, and which themselves appear to be linked to the host genotype.

## Discussion

Life in the phyllosphere is harsh for microorganisms, as they are exposed to a variety of abiotic site-specific environmental stressors including fluctuating temperatures, humidity and UV radiation, in addition to defense mechanisms of the host plant. In grapevine-associated microbial communities, such site-specific effects are the main driver in shaping these communities and likely obscure the secondary effect of the host genotype in microbiome assembly (Griggs *et al*., 2021). Yet, host genotype plays a pivotal role in shaping endophytic microbiota, which becomes more apparent when comparing varieties within specific regions or sites (Bokulich *et al*., 2014, 2016; Portillo *et al*., 2016; Awad *et al*., 2023). Different grapevine cultivars have distinct morphological traits, including cuticle composition, cluster compactness, and leaf morphology, (Gabler *et al*., 2003; Chitwood *et al*., 2014; Tello & Ibáñez, 2018) creating unique microenvironments on their surface. Microbial biofilms often localize around structural features like veins and stomata (Vorholt, 2012), and these physical niches, along with differences in plant surface structures, thereby directly influence microbial community assembly. Furthermore, plants produce secondary metabolites and antimicrobial compounds that have been shown to affect susceptibility to fungal pathogens, such as *B. cinerea* (Gabler *et al*., 2003; Herzog *et al*., 2015; Tello & Ibáñez, 2018). In grapevines, the berry microbiome also directly influences fermentation outcomes and wine characteristics. To our knowledge, no study has yet explored the genomic basis of microbiome assembly in grapevine (Tello & Ibáñez, 2023). Here, we present strong evidence for a robust effect of grapevine variety on microbial community composition. The QTL analysis shed light on the genetic mechanisms affecting microbiome recruitment in grapevines.

Our preliminary study robustly showed that varieties harbor distinct microbial communities, even at a close range, in mono-varietal blocks within a single vineyard with no major topographical differences between them. Fungal communities thereby appeared more spatially heterogeneous than bacterial communities, which is likely due to their larger spore size leading to a greater dispersion limitation (Miura *et al*., 2017). Yet, within the same mono-varietal blocks and within the whole vineyard when the phylogenetic distance between the varieties was taken into account, spatial heterogeneity of microbial communities was negligible. This supports the hypothesis that varieties harbor distinct microbial communities due to some features of their parentage and morphology. Furthermore, it suggests that dispersion, and niche selectivity are important and likely cooperative drivers of microbial community assembly on fruit at the intra-vineyard scale. The differences between blocks could be the result of the selective effects of cluster morphology and exudates, or growth pattern and canopy density, even in blocks trellised the same and in the same row orientation, exerting selective pressures on the microbial communities associated with them. It also could be a possibility that plant-to-plant transmission strengthens this relationship at the block level, if source microbial communities have been shaped by a common niche after primary seasonal seeding.

To further disentangle abiotic from biotic effects on the berry microbiome, we subsequently studied a novel experimental population of 140 F1 genotypes deriving from a cross between Riesling and Cabernet Sauvignon, where intra-vineyard spatial differences among microbial communities were negligible. As the preliminary study showed that variety-specific microbial communities remain temporally stable throughout the ripening stages within a given season we applied two different sampling strategies to elucidate the role of different degrees of ripening versus climatic effects. Microbial communities on grape berry surfaces are known to change throughout the ripening process as the microbiome is reshaped by host-microbe interactions, for example by yeast proliferation due to increased sugar exudates (Griggs *et al*., 2021). The composition of these exudates, which includes sugars, mineral nutrients, and organic acids, also shifts during ripening, influencing the microbial communities on the berry surface (Griggs *et al*., 2021). The climatic environment during harvest season also impacts the berry microbiome, thus applying both approaches has the potential to unravel which associations are robust throughout the ripening process and which influence the genotype exerts only during peak ripeness. We observed significant variation in microbial communities between years, which aligns with findings from previous studies (Bokulich *et al*., 2014; Zarraonaindia *et al*., 2015; Cureau *et al*., 2021) and is unsurprising given the difference in climatic conditions between years, e.g. in accumulated growing degree days. However, we found that particularly fungal communities were modestly, yet robustly associated with the grape berry color in both years. This may be caused by a combination of direct effects, such as the antimicrobial properties of anthocyanins present in red grape varieties, and indirect effects, like physiological changes in the berries associated with the expression of color-related genes. Furthermore, microbial communities of vines from the same genotype were consistently more similar to each other, regardless of the sampling strategy. This reveals, for the first time, the dominant role of the grapevine genotype in shaping microbial communities, even in closely related plants, and irrespective of ripeness and environmental variation. Our analysis also revealed that certain fungal and bacterial communities remain stable across years, indicating that the grapevine host exerts a steady influence on microbial recruitment.

Similarly, we found that many putative fungal pathogens were robustly associated with the host genotype in both years. Many of these putative pathogens are mapping to the same genetic loci, which is also reflected in the network analysis, in which they are closely correlated with each other and commonly form central hubs. Of particular interest is the strong genetic association observed with *Botrytis* spp., an important berry-infecting fungal pathogen that causes fruit deterioration during ripening (Rienth *et al*., 2021). As *Botrytis* spp. induces host cell death, it often co-occurs with plant-matter-decomposing (saprophytic) and endophytic fungi. In line with existing literature, we found co-occurring saprophytic endophytes *Cladosporium* and *Alternaria spp.* (Rienth *et al*., 2021), as well as *Mycosphaerella tassiana* and *Epicoccum nigrum*, some of which are even linked to the same genetic marker as *Botrytis* spp. This co-localization of QTLs for both pathogenic and saprophytic species suggests a potential shared genetic basis for their presence in the grapevine microbiome. Notably, a substantial number of taxa – many of which show strong associations with the host genotype across both years – are endophytes, including *Cladosporium* spp., *M. tassiana*, *E. nigrum*, and *Bacillus* spp. Endophytes are especially interesting for their capacity to inhabit plant tissue without causing visible disease symptoms and for their frequent co-occurrence with pathogens, which raises questions about their role in plant disease dynamics and their interactions with the host (Crandall *et al*., 2020). This observation also aligns with an overall shift in plant pathology: moving from viewing diseases as interactions between a single host and a single pathogen, to recognizing more intricate, multi-species and often multi-strain disease complexes (Dutt *et al*., 2022), including the Esca disease in grapevines (Morales-Cruz *et al*., 2018).

The grape berry microbiome plays a crucial role in wine fermentation and ultimately influences wine characteristics (Garofalo *et al*., 2015; Pinto *et al*., 2015; Bokulich *et al*., 2016). Here, we present the first evidence that host genotype directly influences the recruitment of specific fermentative organisms, such as *Saccharomyces cerevisiae* and *Rhodosporidiobolus* spp. While previous research has documented variety-specific associations with distinct fermentative organisms, including different strains of *S. cerevisiae* (Blanco *et al*., 2006; Cordero-Bueso *et al*., 2011), the genetic mechanisms underlying this specificity have remained elusive due to the inherent complexity of microbial systems. This is also reflected in the central role of *S. cerevisiae* in the network analysis in both study years, where it functions as a connecting node between bacterial and fungal communities in 2020 and showing strong positive correlations with various endophytic fungi in 2022. Notably, the QTL mapping demonstrated that both yeasts are associated with the same genomic hotspot that was also linked to pathogenic organisms. This co-occurrence, for example of *S. cerevisiae* in *Botrytis* infected berries, has been previously reported and suggested to be caused by pathogen-induced berry damage creating favorable conditions for yeast colonization or complex antagonistic or synergistic interactions (Di Canito *et al*., 2021).

QTL interval analyses revealed that a diverse range of microbes, including fermentative yeasts and pathogens, mapped to intervals containing genes associated with plant immune responses. Notably, two of the strongest and robust associations with various pathogenic fungi as well as commensals across both years mapped directly to chitinase-related genes. Chitinase can degrade the fungal cell wall component chitin, and thereby releasing nutrients for other microbes. Strikingly, the chitinase precursor gene within this hotspot also showed robust associations with bacterial genera such as *Lysinibacillus* and *Bacillus*. These bacterial genera are themselves producers of antifungal compounds, including chitinases (Ahsan & Shimizu, 2021). This finding suggests that alleles modulating chitinase activity could influence nutrient cycling and bacterial colonization. Similarly, an ankyrin-repeat protein gene, likely functioning as an immune receptor recognizing specific to fungi (Kolodziej *et al*., 2021), was associated with *Alternaria* spp. and *S. cerevisiae*. Prior studies have demonstrated the overexpression of ankyrin-repeat protein (Vigneron *et al*., 2023) and cysteine synthase genes (Azri *et al*., 2021) in grapevines under stress. Ankyrin-repeat proteins, in particular, have been linked to pathogen resistance, including defense against black rot (Rex *et al*., 2014). Our findings thus underscore the intricate nature of grapevine microbiome assembly and emphasize the need for targeted studies to elucidate the precise mechanisms by which host genotype influences microbial recruitment. We anticipate that our current findings could serve as a roadmap for targeted experiments to investigate and confirm causative mechanisms of microbiome recruitment by the grapevine host. Such studies could include application of synthetic, defined microbial communities, and transcriptomics to uncover the actual host response, as well as targeted gene knockouts in the host.

Overall, we provide evidence that even full-sib grape genotypes harbor distinct microbial communities and give a first insight into potential genetic mechanisms contributing to this genotype-specific recruitment. By combining approaches of quantitative genetics with community-level microbiome data, we establish a framework for understanding the complex interactions between genotype, environment, microbiome, and management practices that shape viticultural ecosystems. Our findings could have important implications for broader research in perennial plants, particularly in breeding programs aiming to enhance natural disease resistance and to further develop more sustainable practices for example by reducing dependency on fungicides. Future research should focus on characterizing the specific causal, molecular mechanisms of host-mediated microbial recruitment, exploring synergistic relationships between beneficial microbes, and assessing their impact on plant health.

## Methods

### Establishing the RxCS Mapping Population

The population used in this study consisted of 138 full-sibling progeny resulting from a cross between *Vitis vinifera* ssp. *vinifera* cv. Riesling clone FPS 24 and *V. v. vinifera* cv. Cabernet Sauvignon clone FPS 08. The cross was performed in 1994 by M.A. Walker and C.P. Meredith and planted at the Wolfskill Germplasm Repository (Riaz *et al*., 2004). In order to have sufficient material for evaluation of viticultural and enological traits, budwood from the original vines was grafted onto certified virus-free *V. rupestris x riparia* Couderc 3309 rootstock in 2017 and planted at the Viticulture and Enology Department Experimental Station (Oakville, Napa, CA) in 2018. The population was planted as a randomized complete block vineyard consisting of three biological replicates of each genotype in three adjacent blocks for a total of nine vines per genotype, including both parental cultivars. Training consisted of bilateral cordons in a modified vertical shoot positioning system pruned to two-bud spurs.

### Genotyping and Map Construction

Genotyping-by-sequencing (GBS) was used to generate markers for high-density linkage mapping. The RxCS population was genotyped using GBS as previously described (Hyma *et al*., 2015). DNA from young leaves was extracted using the DNeasy 96-well DNA extraction kit (Qiagen, Valencia CA, USA). Libraries were prepared using ApeKI as the digestion enzyme and sequenced with HiSeq2000 (Illumina Inc., San Diego CA, USA) at the Institute of Biotechnology, Genomics Facility (Cornell University, Ithaca, NY).

Raw sequencing reads for the RxCS population were analyzed using the Tassel 5 GBS v2 pipeline with default parameters (Glaubitz *et al*., 2014). SNPs were identified by aligning sequence reads against the haplotype 1 of the genome of Cabernet Sauvignon clone 08 (v.1.1) (Massonnet *et al*., 2020) with BWA MEM and default parameters (Li & Durbin, 2009; Chin *et al*., 2016). The Tassel 5 standalone program was used to detect SNPs and filter SNPs for coverage depth (minimum read depth > 6) and frequency (minor allele frequency > 0.1). SNPs and genotypes with more than 10% missing data were removed, and SNPs within 64 base pairs were merged. A high-density genetic linkage map was constructed as previously outlined (Lopez-Moreno *et al*., 2023) using ABHGenotypeR for imputation of missing marker data and the Kosambi mapping function from the ASMap package for map construction (Taylor & Butler, 2017; Furuta *et al*., 2017).

### Berry Sampling and DNA extraction

For the preliminary monovarietal block vineyard study, grape berries were collected at ripeness before harvest from a teaching vineyard in Davis, California (38.52675, −121.78946) in 2013. This vineyard is planted with various varieties, with all of them cordon-trained with vertical shoot positioning and farmed conventionally under the same management regime. Samples of five varieties including Chardonnay, Cabernet Sauvignon, Grenache blanc, Grenache noir, and French Colombard, planted in mono-variety blocks within this single vineyard were sampled across the breadth of each block 8 vines in three rows each. Berries were aseptically collected at commercial ripeness from a single cluster in sterile Whirl-Pak sampling bags (Weber Scientific, Hamilton, NJ, USA). Upon collection, berry samples were crushed within the sterile bags immediately upon return to the laboratory, and frozen at −80°C until processing. For DNA extraction, crushed berry samples were thawed and centrifuged as must at 4,000 x g for 15 min, washed in ice-cold PBS, and suspended in 200uL DNeasy lysis buffer, with an added 40mg/mL lysozyme, and then incubated at 37°C for 30 min. Samples from fruit swabs were thawed and aseptically transferred to ZR-96 Fecal DNA MiniPrep Kit (Zymo Research, Irvine, CA) tubes. All samples were then processed according to the manufacturer’s protocol, with the addition of a 2 min at maximum speed bead beater cell lysis step using a FastPrep-24 bead beater (MP Bio, Solon, OH).

For the F1 population, grape samples were collected from the RxCS experimental population planted at the UC Davis research station located in Oakville, Napa County, California (38.43670, −122.40353). This experimental vineyard with F1 progeny of Riesling and Cabernet Sauvignon is planted in a randomized complete block design (140 genotypes including parents planted in triplicate in 3 blocks = 1260 vines, with 9 vines per genotype). In 2020, samples of each genotype and each block (140 genotypes x 3 blocks = 420 samples) were collected at equivalent ripeness (24 °Brix) over a time period of almost two months (August 8th 2020 until September 26th 2020), while in 2022, samples were collected within 2 days (02-09-2022 and 03-09-2022). These samples were transported back to the lab on ice, and stored at −80 °C until processing. Grapes were thawed and pressed in a stomacher. Thereof 250 µL were used for DNA extraction with the MagAttract PowerSoil Pro DNA Kit (QIAGEN, cat. no. 47109) and PowerBead Pro Plate (QIAGEN, cat. no. 19311) according to the manufacturer’s instructions. Cell lysis was performed on a Bead Mill (Retsch) and extraction on a KingFisher Flex (Thermo Fisher Scientific). Extracted DNA was stored at −20°C until further processing.

### Library Preparation and Sequencing

For the monovarietal block study, the amplification and sequencing were completed as previously described (Griggs *et al*., 2025). Briefly, the V4 domain of the 16S rRNA gene was amplified using the universal primer pair F515 (5’-GTGCCAGCMGCCGCGGTAA-3’) and R806 (5’-GGACTACHVGGGTWTCTAAT-3’) (Caporaso et al. 2011). The internal transcribed spacer 1 (ITS1) loci were amplified with the BITS (5′–ACCTGCGGARGGATCA–3′) and B58S3 (5′–GAGATCCRTTGYTRAAAGTT–3′) primer pair (Bokulich and Mills 2013). Extracted DNA was checked for quality with gel electrophoresis and a NanoDrop spectrophotometer (Thermo Fischer Scientific, DE, USA). Amplicons of the V4 and ITS regions were separately pooled in equimolar ratios, purified using the Qiaquick spin kit (Qiagen), and submitted to the UC Davis Genome Center DNA Technologies Core Facility (Davis, Ca, USA) for Illumina paired-end library preparation, and 250-bp paired-end sequencing on an Illumina MiSeq (Illumina, Inc. CA, USA).

For the RxCS project, amplicon sequencing was done using the HighALPS ultra-high throughput library preparation protocol based on a unique dual index (UDI) strategy with custom 12 nt long Golay barcodes (Flörl *et al*., 2024). To profile present bacterial communities, we performed amplicon sequencing of the hypervariable V4 region of the 16S rRNA gene using the updated 515F primer (5’-GTGYCAGCMGCCGCGGTAA-3’) (Parada *et al*., 2016) and 806R primer (5’-GGACTACNVGGGTWTCTAAT-3’) (Apprill *et al*., 2015). To amplify the low microbial biomass samples, we performed a nested PCR. In the first PCR, all template DNA was enriched with 515F/806R primers, and in the second PCR the amplicons were UDI barcoded as well as plant host plastid and mitochondrial 16S rRNA gene contaminations depleted with peptide nucleic acid (PNA) PCR clamps (Flörl & Bokulich, 2024). Therefore, we first set up a 25 µL PCR reaction with 12.5 µL of 2x KAPA HiFi HotStart ReadyMix (Roche, Cat. No. 07958935001), 0.5 µM of each primer (Microsynth) and 2.5 µL of extracted DNA. All PCR reactions were set up using an epMotion liquid handling platform. Reaction conditions were initially 95°C for 5 min, followed by 20 cycles of 95°C for 30 sec, 55 °C for 30 sec, and 72°C for 30 sec, and a final extension of 72°C for 5 min. From this enriched template we used 1 µL per 25 µL reaction for a second PCR with 12.5 µL of 2x KAPA HiFi HotStart ReadyMix, 0.5 µM of UDI primer combinations as well as 0.5 µM or mPNA and pPNA clamps. The PNAs were vortexed and incubated at 60°C for 10 min before use. The subsequent reaction conditions were 95°C for 5 min, followed by 25 cycles of 95°C for 30 sec, 78°C for 5 sec for PNA annealing, 50°C for 30 sec, and 72°C for 30 sec, and a final extension of 72°C for 5 min. Notably, the presence of PNAs in the reaction showed that a lower annealing temperature increased yield. For fungal communities, we amplified the first loci in the internal transcribed spacer (ITS) region with above described BITS and B58S3 primers (Bokulich & Mills, 2013). For the library amplification, we set up 25 µL PCR reactions with 12.5 µL of 2x KAPA HiFi HotStart ReadyMix, 0.5 µM of each BITS/B58S3 primer, and 2.5 µL of extracted DNA. Reaction conditions consisted of initial 95°C for 5 min, followed by 34 cycles of 95°C for 30 sec, 49°C for 30 sec, and 72°C for 30 sec, and a final extension of 72°C for 5 min. Thereof we used 1 µL template DNA to barcode the amplicons with 12.5 µL of 2x KAPA HiFi HotStart ReadyMix and 0.5 µM UDI primer combinations. The barcoding PCR conditions consisted of initial 95°C for 5 min, followed by 8 cycles of 95°C for 30 sec, 49°C for 30 sec, and 72°C for 30 sec, and a final extension of 72°C for 5 min.

All amplicons were purified with Agencourt AMPure XP magnetic beads (Beckman, Cat No. A63882) with a ratio of 0.7x to remove primer dimers and small fragments on the KingFisher Apex. Resulting DNA concentrations were measured in duplicates with Qubit dsDNA High Sensitivity Assay (Thermo Fisher Scientific, Cat No. Q32854) on a Tecan Spark Microplate Reader. Amplicons were pooled equimolarity in two separate pools with a liquid handling platform (Brand GmbH, Wertheim, Germany) and quality controlled on a Tapestation (Agilent). To increase yield we reconditioned each pool separately with 12.5 µL of 2x KAPA HiFi HotStart ReadyMix, 0.4 µM of the standard Illumina P5 (5′-AATGATACGGCGACCACCGAGATCT-3′) and P7 (5′-CAAGCAGAAGACGGCATACGAGAT-3′) PCR primers, 2 µL template of the bacterial pool, and 3 µL template of the fungal pool. Reconditioning PCR conditions were set to initial 95°C for 3 min, followed by 4 cycles of 98°C for 20 sec, 62°C for 15 sec, and 72°C for 30 sec, and a final extension of 72°C for 1 min. Ultimately we conducted a two-sided clean up with 0.2x magnetic bead ratio, followed by 0.7x magnetic bead ratio of the supernatant to remove remaining genomic DNA as well as primers from the reconditioning. The pools were combined and submitted to the Functional Genomics Center Zürich for Paired End 250 bp sequencing on the Illumina NovaSeq 6000 SP 500 cycles with 20 %PhiX. In total we obtained 552’726’447 reads (16S: 281’134’325, ITS: 271’592’122).

### Microbiome Diversity Analysis

For the preliminary monovarietal block study, raw Illumina fastq files were demultiplexed and barcodes were stripped with Sabre (UC Davis Bioinformatics, 2011). Sequences were then imported into QIIME2 version 2018.6 (Bolyen *et al*., 2019) and before being denoised with DADA2 (Callahan *et al*., 2016) to generate ASVs (amplicon sequence variants). Briefly, V4 forward reads were truncated at 240 nucleotides based on quality scores; primers were removed by trimming. Forward and reverse reads were then merged, and chimeras removed using DADA2. Only forward reads were used in the ITS data due to low-quality reverse reads, and truncated at 242 nucleotides due to low-quality reverse reads, and ASVs otherwise processed as described for bacterial reads. Contaminations were removed with decontam Daivis 2018 (Davis *et al*., 2017) (via the q2-quality-control plugin) against the blanks, further we removed singletons and features not classified to phylum level. Additionally we filtered non-target reads, i.e. mitochondria and chloroplasts from the 16S rRNA gene data and macrofungal taxa from the ITS data. For the microbial diversity, we rarefied to 1,000 reads per sample for bacteria and fungi and calculated non-phylogenetic alpha and beta diversity metrics with q2-diversity. The PERMANOVA was performed with ADONIS permutation-based test (Anderson, 2001; Oksanen *et al*., 2001) (via q2-diversity), mantel tests with q2-diversity and partial mantel tests with vegan (Oksanen *et al*., 2001) (via q2-diversity), all with 999 permutations, respectively.

The microbial diversity analysis for the RxCS dataset was performed in QIIME 2 version 2024.2 (Bolyen *et al*., 2019). Demultiplexed Illumina fastq files were imported, potential adapter remnants trimmed with cutadapt (Martin, 2011) and denoised with DADA2 (Callahan *et al*., 2016) as paired-end reads for 16S and single-end reads for ITS. Resulting ASVs were taxonomically classified with the q2-feature-classifier plugin (Bokulich *et al*., 2018), in which we used a naïve Bayes taxonomy classifier trained on the 99 % SILVA (Quast *et al*., 2012) 16S rRNA gene database (138 release), trimmed to the 515F-806R (V4) region. Similarly, fungal ASVs were classified with the UNITE database (v9.0, Version 18.07.2023) (Abarenkov *et al*., 2023). Furthermore, we removed contaminations with decontam from the DNA extractions negative controls. We filtered ASVs from non-target reads, namely fruiting body forming mushrooms from ITS and host DNA from 16S, as well as all ASVs not classified to at least phylum level. Additionally, we filtered samples with less than 10 associated ASVs, as well as samples that had started to ferment (i.e. with >40% of *S. cerevisiae*). For the microbial diversity we rarefied to 1,000 reads per sample for bacteria and 80,000 reads per sample for fungi. Diversity analysis was performed with the same packages and tools described for the preliminary study above. For the network analysis, FeatureTables were analyzed separately by year and collapsed per genotype with FlashWeave (Tackmann *et al*., 2019) via q2-makarsa (Kaehler, Benjamin, 2022).

### Grapevine Variety Phylogeny Analysis

Genomic reads were quality-filtered and trimmed using Trimmomatic (v. 0.36, “ILLUMINACLIP 2:30:10, LEADING 7, TRAILING:7, SLIDINGWINDOW:10:20, MINLEN:36) (Bolger *et al*., 2014). Filtered read pairs were mapped using BWA MEM (v. 0.7.12-r1039, default parameters) (Li, 2013) using *Vitis vinifera* PN40024 genome V2 as reference (Canaguier et al. 2017). Alignments were randomly downsampled to represent a maximum read coverage of 40X using Samtools (v. 1.7) (Li *et al*., 2009). Variants were called separately for each sample using GATK HaplotypeCaller (v. 4.0.12.0, “--base-quality-score-threshold 20 -- sample-ploidy 2”) (Poplin *et al*., 2017) The gVCF were then combined with GATK CombineGVCFs (v. 4.1.4.1-83-g031c407-SNAPSHOT) (Poplin *et al*., 2017) and genotyped with GATK GenotypeGVCFs (v. 4.1.4.1-83-g031c407-SNAPSHOT, “--sample-ploidy 2 --max-alternate-alleles 12”) (Poplin *et al*., 2017). The resulting dataset was filtered removing a) sites not genotyped in all samples, b) repetitive regions, c) non-variant sites, d) variants detected on the undermined chromosome (chr00). The filtered variant set was converted from VCF to PHYLIP format using vcf2phylip (v. 2.0) (Ortiz, 2019) The phylogenetic tree was inferred with a 100 iteration boostrap procedure using RAxML-NG (v. 0.9.0 released on 20.05.2019) with the option “--seed 12345 --tree pars{10} --bs-trees 100 --model GTR+G”) (Kozlov *et al*., 2019). Graphically representation of the phylogenetic tree was produced using FigTree (v. 1.4.4) (https://github.com/rambaut/figtree/). Pairwise distance calculation was performed using PLINK (v. 1.90b5.2) and the parameter “--distance square 1-ibs” (Chang et al. 2015) to obtain the Hamming distance expressed as reciprocal of the identity-by-state values.

### QTL Mapping

We performed QTL mapping using the Cabernet Sauvignon Haplotype 1 genetic map, which contains 5’386 with a density of 28.96 ± 22.25 cM and a genotype completion of 97.8%. The quality of the genetic map was checked with r/qtl (Broman & Sen, 2009). We performed mapping of the microbial features using ASVs clustered at 99% similarity (OTUs) to collapse variants and reduce noise. The years were analyzed separately, and at different taxonomic levels: genus, family and order level respectively. We stringently filtered out resulting OTUs which occurred in less than ten samples, and less than five samples at higher taxonomic levels. Ultimately, we calculated the median read count per genotype before converting the FeatureTable to a presence/absence matrix. This was then used for interval mapping with r/qtl (Broman & Sen, 2009) against the Cabernet Sauvignon Haplotype 1 genetic map using a Haley-Knott regression without inserting pseudo-markers. The LOD threshold to assess significance was calculated for each phenotype (i.e. microbial feature) individually with 1,000 permutations. QTLs were considered significant with a LOD above threshold as well as a genome-wide p-value < 0.05. Furthermore, we calculated the LOD support interval and Bayes credible interval to estimate the QTL interval, as well as the effect size with r/qtl. Within the respective BCI we parsed genes and functional annotation from the *Vitis vinifera* cv. Cabernet Sauvignon cl. 08 v1.1 reference genome (Cochetel & Cantu, 2024). Retrieved genes were also matched with the *Vitis vinifera* PN40024 reference genome (v1), from which we retrieved Arabidopsis homologs which were then researched in the Arabidopsis Information Resource (TAIR) database (Reiser *et al*., 2024). Gene Ontology (GO) enrichment analysis was performed using the python library GOATOOLS (Klopfenstein *et al*., 2018) using the annotated *Vitis vinifera* cv. Cabernet Sauvignon cl. 08 v1.1 reference genome as background. All plots were generated using matplotlib (The Matplotlib Development Team, 2024) or seaborn (Waskom, 2021).

## Data Availability

Amplicon sequencing data is available from the Sequence Read Archive (SRA) under accession number PRJEB85265 (16S) and PRJEB85263 (ITS).

## Code Availability

All code used in this study is available on GitHub in the following repositories: (i) for the preliminary study (https://github.com/LenaFloerl/Tyree-genotype.git), (ii) for the microbiome analysis of the RxCS population (https://github.com/LenaFloerl/RxCS.git) and (iii) of the QTL mapping (https://github.com/LenaFloerl/RxCS_mapping.git).

## Acknowledgement

The authors gratefully acknowledge financial support from the Swiss National Science Foundation [Grant Number: 310030_204275] (to NAB) and the Swiss Government Excellence Ph.D. Scholarship (to LF). The project was also partially supported by the Ray Rossi Endowment in Viticulture and Enology (to DC) and the E. & J. Gallo Winery.

We thank Bruce McDonald (ETH Zurich) and Karl Broman (University of Wisconsin-Madison) for their advice and input on QTL mapping. The authors thank Serafina Plüss and the Genetic Diversity Centre (GDC) of ETH Zurich for supporting the library preparation. The microbiome amplicon sequencing was performed at the Functional Genomics Center Zurich (FGCZ) of University of Zurich and ETH Zurich.

We thank Andrea Minio (UC Davis) for bioinformatic support, Guillermo Garcia Zamora (UC Davis) for managing the experimental vineyards, and Drs. Lance Cadle Davidson (USDA ARS) and Bruce Reisch (Cornell University) for assistance in generating the GBS data as part of the U.S. Department of Agriculture-Specialty Crops Research Initiative grant number 2011-51181-30635.

## Author Contributions

The study was conceived by NAB and DC and supervised by NAB. LF performed the microbiome analysis, QTL mapping, and all statistical analyses, and wrote the article with review and contributions from NAB, RG, DC, and JL. RG collected samples and planned and performed analysis of the monovarietal vineyard. JL and the research group of DC (MM, NC, RFB) established the experimental vineyard and the genetic maps, collected samples, and prepared the grapevine phylogeny.

## Competing Interests

The authors declare no competing interests.

## Notes

### Competing Interest Statement

The authors have declared no competing interest.

