## Supporting Information for "The Genetic Basis of Microbiome Recruitment in Grapevine and its Association with Fermentative and Pathogenic Taxa"

### 1. Supplementary Figures

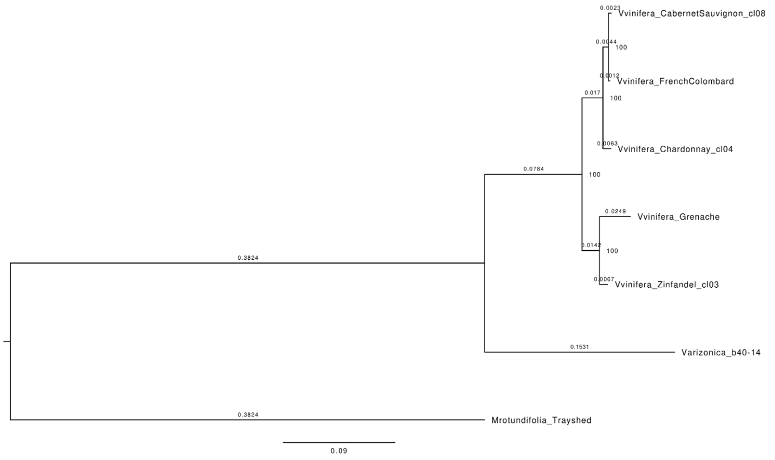


**Supplementary Figure 1.** Phylogenetic tree of grapevine varieties used in the partial Mantel test.

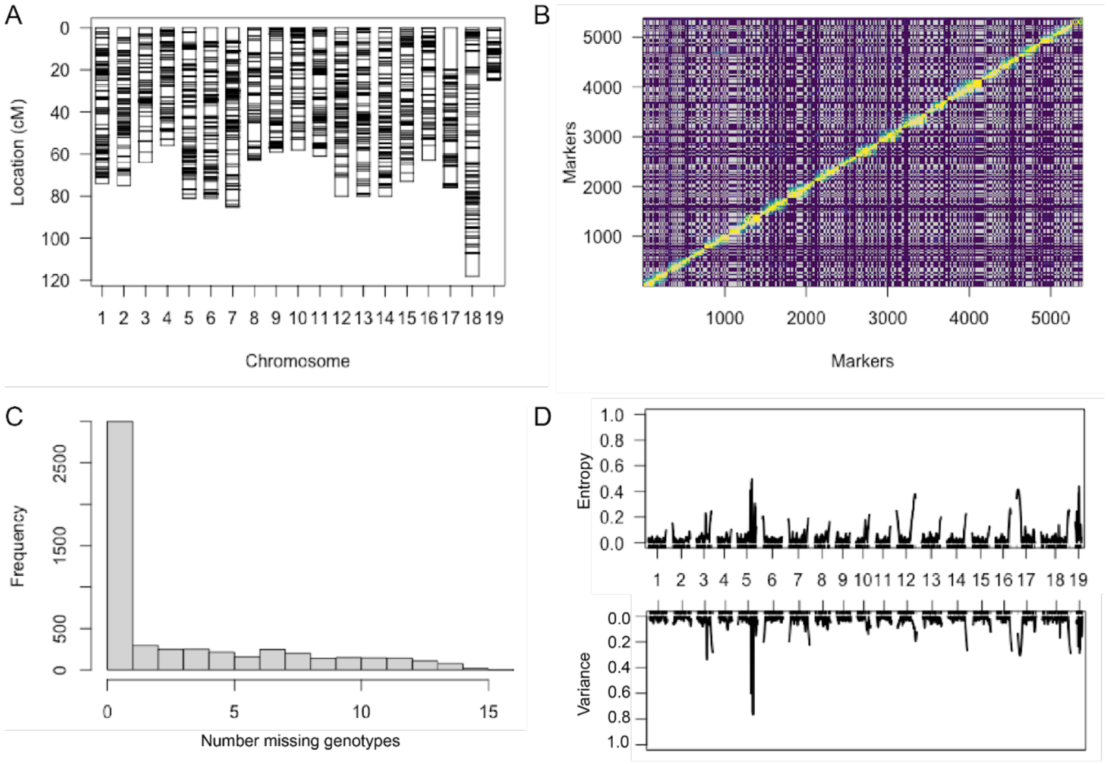


**Supplementary Figure 2.** Quality control of the genetic maps. A. The genetic map used in the QTL mapping has an even distribution of markers across the 19 chromosomes. B. Plot of the estimated recombination fractions between markers shows linkage between markers and confirms their correct order. C. Overall our genetic maps have a high quality as most markers are typed for all genotypes. D. To reveal potentially missing genotyping information we calculate the entropy and variance of each marker.

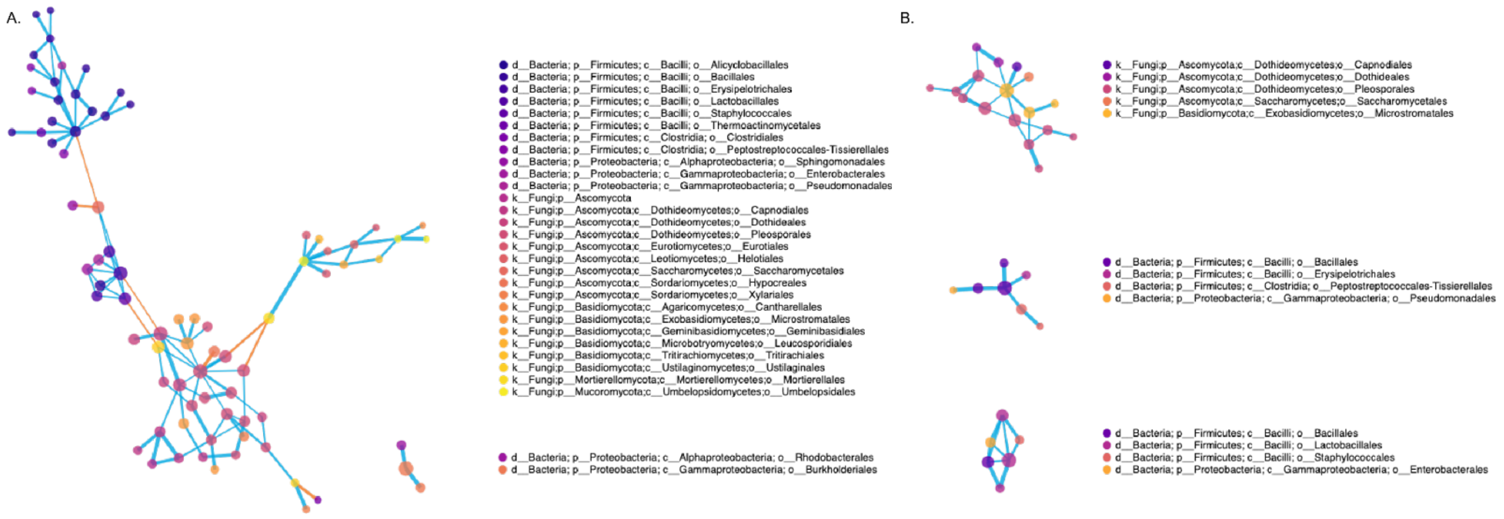


**Supplementary Figure 3.** Complementary network visualization to Figure 5 showing the communities of fungal and bacterial features for 2020 (A) and 2022 (B) with the nodes colored by taxonomic assignment on the order level.

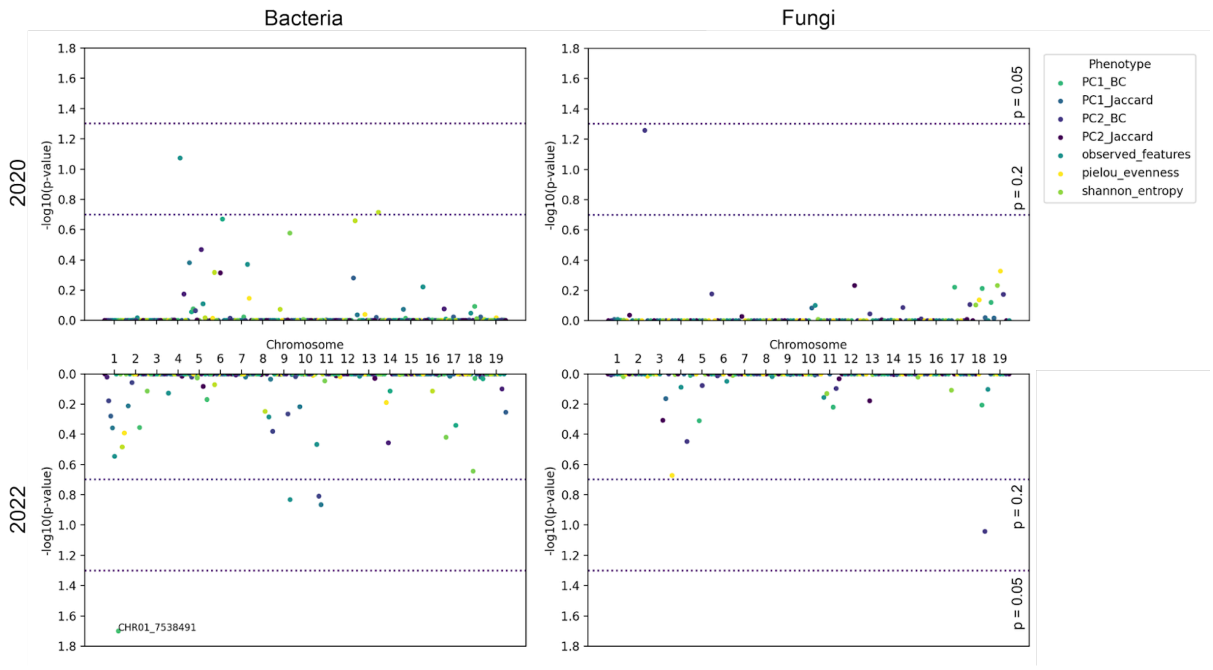


**Supplementary Figure 4.** Manhattan plots of the mapped diversity metrics for bacteria (left) and fungi (right) of 2020 (top) and 2022 (bottom). The marker with significant QTLs on chromosome 1 associated for bacteria in 2022.

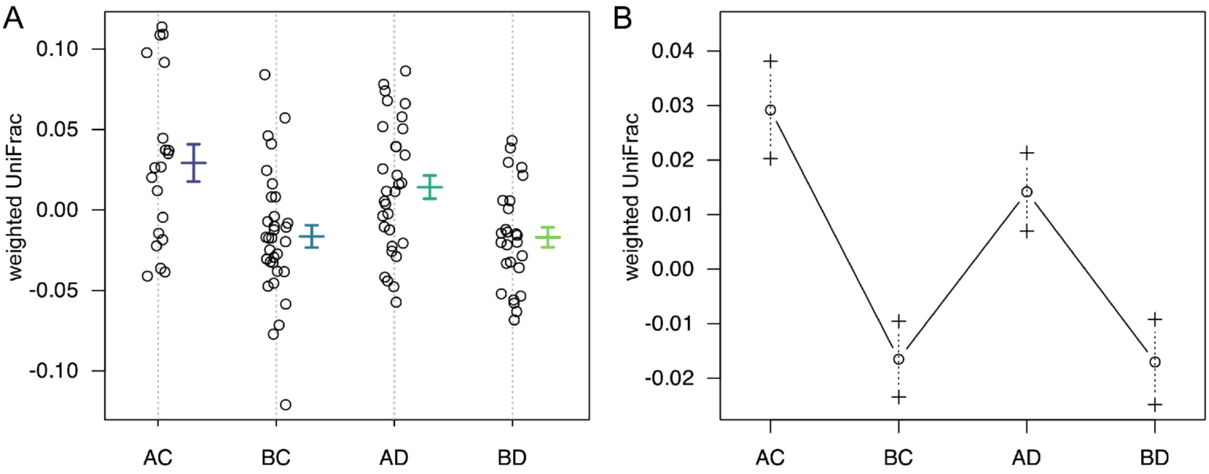


**Supplementary Figure 5.** (A.) Phenotypic variation for the weighted UniFrac metric of bacterial communities in 2022 over the alleles at the significant QTL of CHR01_7538491 and (B.) the estimated effects of these genotypes (Error bars are ± 1SE).

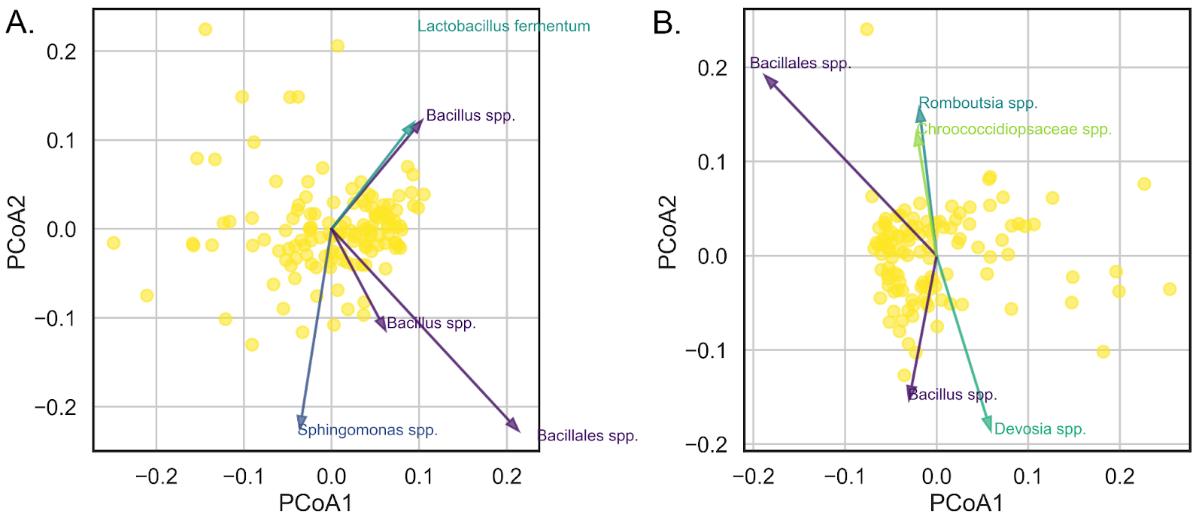


**Supplementary Figure 6.** Biplots showing distribution of samples according to the weighted UniFrac metric of bacterial communities in 2020 (A) and 2022 (B) and the top 5 OTUs contributing to the spread thereof.

### 2. Supplementary Tables

**Supplementary Table 1.** Temperature data [°F] and accumulated growing degree days (GDD) for the two sampling periods in 2020 and 2022.

| | | Date | Accumulated GDD | Mean Temperature | Max. Temperature | Min. Temperature |
|---|---|---|---|---|---|---|
| **2020** | **Start Sampling** | 2020-08-08 | 2001.09 | 69.5 | 85.0 | 54.0 |
| | **End Sampling** | 2020-09-26 | 2930.68 | 74.0 | 93.0 | 55.0 |
| **2022** | **Start Sampling** | 2022-09-02 | 2474.71 | 68.5 | 89.0 | 48.0 |
| | **End Sampling** | 2022-09-03 | 2499.71 | 75.0 | 98.0 | 52.0 |

**Supplementary Table 2.** Comparing beta diversity distances across all vines (All), between different genotypes (Between) and within the same genotype (Within) using the Kruskal-Wallis test reveals that diversity distances are significantly more similar among vines of the same genotype (considering significance with a p-value < 0.05).

| | | | Bray Curtis | Jaccard | weighted UniFrac | unweighted UniFrac |
|---|---|---|---|---|---|---|
| **Bacteria** | **2020** | All : Within | 2.12E-07 | 2.70E-12 | 3.60E-03 | 1.46E-11 |
| | | Between : Within | 1.84E-07 | 2.10E-12 | 3.43E-03 | 1.16E-11 |
| | **2022** | All : Within | 1.13E-03 | 2.98E-03 | 6.77E-07 | 3.49E-05 |
| | | Between : Within | 1.07E-03 | 2.83E-03 | 5.92E-07 | 3.18E-05 |
| **Fungi** | **2020** | All : Within | 4.19E-08 | 4.85E-14 | | |
| | | Between : Within | 3.59E-08 | 3.63E-14 | | |
| | **2022** | All : Within | 2.43E-05 | 1.42E-02 | | |
| | | Between : Within | 2.20E-05 | 1.37E-02 | | |

**Supplementary Table 3.** Overview of the number of mapped features per year and marker gene as well as the resulting significant QTLs.

| | | 2020 | | | | | 2022 | | | | |
|---|---|---|---|---|---|---|---|---|---|---|---|
| | | cASVs | Genera | Family | Order | Total | cASVs | Genera | Family | Order | Total |
| **Fungi** | **mapped features** | 3944 | 484 | 254 | 112 | **4794** | 513 | 148 | 101 | 56 | **818** |
| | **sign. QTLs** | 47 | 8 | 6 | 4 | **65** | 27 | 8 | 6 | 3 | **44** |
| **Bacteria** | **mapped features** | 2731 | 825 | 368 | 218 | **4142** | 410 | 258 | 150 | 97 | **915** |
| | **sign. QTLs** | 22 | 14 | 13 | 8 | **57** | 10 | 8 | 5 | 3 | **26** |

**Supplementary Table 4.** All markers at which a significant QTL was detected over the different datasets (in blue) reveal that some are conserved across years or between bacteria and fungi (highlighted in bold).

| | Fungi 2020 | | | | Fungi 2022 | | | | Bacteria 2020 | | | | Bacteria 2022 | | | |
|---|---|---|---|---|---|---|---|---|---|---|---|---|---|---|---|---|
| marker | OTUs | genera | family | order | OTUs | genera | family | order | OTUs | genera | family | order | OTUs | genera | family | order |
| **CHR05_24281346** | 26 | 0 | 0 | 0 | 0 | 1 | 1 | 0 | 2 | 0 | 0 | 0 | 1 | 1 | 2 | 1 |
| **CHR05_24608659** | 6 | 0 | 0 | 0 | 22 | 3 | 1 | 1 | 0 | 0 | 0 | 0 | 1 | 0 | 0 | 0 |
| **CHR05_15679446** | 3 | 0 | 0 | 0 | 0 | 0 | 0 | 0 | 3 | 0 | 0 | 0 | 0 | 0 | 0 | 0 |
| CHR01_4055584 | 1 | 1 | 0 | 0 | 0 | 0 | 0 | 0 | 0 | 0 | 0 | 0 | 0 | 0 | 0 | 0 |
| **CHR05_25035791** | 1 | 1 | 0 | 0 | 1 | 0 | 1 | 0 | 1 | 0 | 1 | 0 | 2 | 0 | 0 | 0 |
| CHR01_533750 | 1 | 0 | 0 | 0 | 0 | 0 | 0 | 0 | 0 | 0 | 0 | 0 | 0 | 0 | 0 | 0 |
| CHR05_19753718 | 1 | 0 | 0 | 0 | 0 | 0 | 0 | 0 | 0 | 0 | 0 | 0 | 0 | 0 | 0 | 0 |
| **CHR05_22428041** | 1 | 0 | 0 | 0 | 1 | 0 | 0 | 0 | 6 | 2 | 1 | 1 | 0 | 0 | 0 | 0 |
| CHR05_24293193 | 1 | 0 | 0 | 0 | 0 | 0 | 0 | 0 | 0 | 0 | 0 | 0 | 0 | 0 | 0 | 0 |
| **CHR05_24493215** | 1 | 0 | 0 | 0 | 0 | 0 | 0 | 0 | 2 | 2 | 1 | 1 | 2 | 1 | 1 | 1 |
| **CHR08_18287104** | 1 | 0 | 0 | 0 | 0 | 0 | 0 | 0 | 0 | 0 | 0 | 0 | 0 | 1 | 1 | 1 |
| CHR11_17209798 | 1 | 0 | 0 | 0 | 0 | 0 | 0 | 0 | 0 | 0 | 0 | 0 | 0 | 0 | 0 | 0 |
| CHR11_6045134 | 1 | 0 | 0 | 0 | 0 | 0 | 0 | 0 | 0 | 0 | 0 | 0 | 0 | 0 | 0 | 0 |
| CHR17_11666180 | 1 | 0 | 0 | 0 | 0 | 0 | 0 | 0 | 0 | 0 | 0 | 0 | 0 | 0 | 0 | 0 |
| CHR19_17331846 | 1 | 0 | 0 | 0 | 0 | 0 | 0 | 0 | 0 | 0 | 0 | 0 | 0 | 0 | 0 | 0 |
| CHR13_6804811 | 0 | 1 | 1 | 1 | 0 | 0 | 0 | 0 | 0 | 0 | 0 | 0 | 0 | 0 | 0 | 0 |
| CHR16_21740082 | 0 | 1 | 1 | 1 | 0 | 0 | 0 | 0 | 0 | 0 | 0 | 0 | 0 | 0 | 0 | 0 |
| CHR16_6298914 | 0 | 1 | 1 | 1 | 0 | 0 | 0 | 0 | 0 | 0 | 0 | 0 | 0 | 0 | 0 | 0 |
| **CHR01_19510296** | 0 | 1 | 1 | 0 | 0 | 0 | 0 | 0 | 0 | 1 | 1 | 0 | 0 | 0 | 0 | 0 |
| **CHR05_19397808** | 0 | 1 | 0 | 0 | 1 | 0 | 1 | 1 | 2 | 1 | 0 | 0 | 1 | 1 | 1 | 0 |
| CHR10_12072879 | 0 | 1 | 0 | 0 | 0 | 0 | 0 | 0 | 0 | 0 | 0 | 0 | 0 | 0 | 0 | 0 |
| CHR12_8475019 | 0 | 0 | 1 | 0 | 0 | 0 | 0 | 0 | 0 | 0 | 0 | 0 | 0 | 0 | 0 | 0 |
| CHR17_14175795 | 0 | 0 | 1 | 0 | 0 | 0 | 0 | 0 | 0 | 0 | 0 | 0 | 0 | 0 | 0 | 0 |
| CHR13_9310000 | 0 | 0 | 0 | 1 | 0 | 0 | 0 | 0 | 0 | 0 | 0 | 0 | 0 | 0 | 0 | 0 |
| CHR19_7737623 | 0 | 0 | 0 | 0 | 1 | 1 | 1 | 0 | 0 | 0 | 0 | 0 | 0 | 0 | 0 | 0 |
| CHR11_11330064 | 0 | 0 | 0 | 0 | 1 | 0 | 0 | 0 | 0 | 0 | 0 | 0 | 0 | 0 | 0 | 0 |
| CHR05_19702836 | 0 | 0 | 0 | 0 | 0 | 1 | 1 | 1 | 0 | 0 | 0 | 0 | 0 | 0 | 0 | 0 |
| **CHR05_24261678** | 0 | 0 | 0 | 0 | 0 | 1 | 0 | 0 | 0 | 0 | 0 | 0 | 1 | 0 | 0 | 0 |
| CHR15_12712252 | 0 | 0 | 0 | 0 | 0 | 1 | 0 | 0 | 0 | 1 | 0 | 0 | 0 | 0 | 0 | 0 |
| CHR15_19813021 | 0 | 0 | 0 | 0 | 0 | 0 | 0 | 0 | 1 | 1 | 0 | 0 | 0 | 0 | 0 | 0 |
| CHR11_15465396 | 0 | 0 | 0 | 0 | 0 | 0 | 0 | 0 | 1 | 0 | 0 | 0 | 0 | 0 | 0 | 0 |
| CHR12_625245 | 0 | 0 | 0 | 0 | 0 | 0 | 0 | 0 | 1 | 0 | 0 | 0 | 0 | 0 | 0 | 0 |
| CHR14_16224251 | 0 | 0 | 0 | 0 | 0 | 0 | 0 | 0 | 1 | 0 | 0 | 0 | 0 | 0 | 0 | 0 |
| CHR17_134705 | 0 | 0 | 0 | 0 | 0 | 0 | 0 | 0 | 1 | 0 | 0 | 0 | 0 | 0 | 0 | 0 |
| CHR18_4958495 | 0 | 0 | 0 | 0 | 0 | 0 | 0 | 0 | 1 | 0 | 0 | 0 | 0 | 0 | 0 | 0 |
| CHR07_19587951 | 0 | 0 | 0 | 0 | 0 | 0 | 0 | 0 | 0 | 1 | 1 | 1 | 0 | 0 | 0 | 0 |
| CHR10_5571895 | 0 | 0 | 0 | 0 | 0 | 0 | 0 | 0 | 0 | 1 | 1 | 1 | 0 | 0 | 0 | 0 |
| CHR12_3309217 | 0 | 0 | 0 | 0 | 0 | 0 | 0 | 0 | 0 | 1 | 1 | 1 | 0 | 0 | 0 | 0 |
| CHR14_3209375 | 0 | 0 | 0 | 0 | 0 | 0 | 0 | 0 | 0 | 1 | 1 | 0 | 0 | 0 | 0 | 0 |
| CHR02_722134 | 0 | 0 | 0 | 0 | 0 | 0 | 0 | 0 | 0 | 1 | 0 | 0 | 0 | 0 | 0 | 0 |
| CHR09_4032132 | 0 | 0 | 0 | 0 | 0 | 0 | 0 | 0 | 0 | 1 | 0 | 0 | 0 | 0 | 0 | 0 |
| CHR02_5830108 | 0 | 0 | 0 | 0 | 0 | 0 | 0 | 0 | 0 | 0 | 1 | 1 | 0 | 0 | 0 | 0 |
| CHR09_3976367 | 0 | 0 | 0 | 0 | 0 | 0 | 0 | 0 | 0 | 0 | 1 | 1 | 0 | 0 | 0 | 0 |
| CHR11_12962073 | 0 | 0 | 0 | 0 | 0 | 0 | 0 | 0 | 0 | 0 | 1 | 1 | 0 | 0 | 0 | 0 |
| CHR03_12211969 | 0 | 0 | 0 | 0 | 0 | 0 | 0 | 0 | 0 | 0 | 1 | 0 | 0 | 0 | 0 | 0 |
| CHR19_3604425 | 0 | 0 | 0 | 0 | 0 | 0 | 0 | 0 | 0 | 0 | 1 | 0 | 0 | 0 | 0 | 0 |
| CHR14_21558244 | 0 | 0 | 0 | 0 | 0 | 0 | 0 | 0 | 0 | 0 | 0 | 0 | 1 | 0 | 0 | 0 |
| CHR15_1432577 | 0 | 0 | 0 | 0 | 0 | 0 | 0 | 0 | 0 | 0 | 0 | 0 | 1 | 0 | 0 | 0 |
| CHR01_607250 | 0 | 0 | 0 | 0 | 0 | 0 | 0 | 0 | 0 | 0 | 0 | 0 | 0 | 1 | 0 | 0 |
| CHR07_24915217 | 0 | 0 | 0 | 0 | 0 | 0 | 0 | 0 | 0 | 0 | 0 | 0 | 0 | 1 | 0 | 0 |
| CHR14_6027109 | 0 | 0 | 0 | 0 | 0 | 0 | 0 | 0 | 0 | 0 | 0 | 0 | 0 | 1 | 0 | 0 |
| CHR17_1853322 | 0 | 0 | 0 | 0 | 0 | 0 | 0 | 0 | 0 | 0 | 0 | 0 | 0 | 1 | 0 | 0 |

**Supplementary Table 5.** Mapping various alpha and beta diversity metrics yielded multiple suggestive QTLs and one significant one for bacteria in 2022 (highlighted in bold) with overall larger support intervals.

| | phenotype | marker | LOD | p-val | BCI [cM] |
|---|---|---|---|---|---|
| **Bacteria 2020** | PC2_wUF | CHR04_5715673 | 4.2 | 0.084 | 13.6 |
| | PC2_uwUF | CHR13_28627421 | 3.8 | 0.193 | 80.0 |
| **Bacteria 2022** | **PC2_wUF** | **CHR01_7538491** | **4.9** | **0.020** | **16.9** |
| | PC1_BC | CHR09_15403002 | 4.1 | 0.148 | 24.5 |
| | PC1_Jaccard | CHR11_569019 | 4.0 | 0.156 | 20.8 |
| | PC1_uwUF | CHR11_569019 | 4.0 | 0.137 | 61.0 |
| **Fungi 2020** | PC2_BC | CHR02_9023323 | 4.4 | 0.055 | 75.0 |
| **Fungi 2022** | pielou_evenness | CHR18_18864631 | 4.3 | 0.091 | 118.0 |

**Supplementary Table 6.** Gene Ontology (GO) enrichment analysis of the genes and corresponding predicted functions from the support intervals retrieved of the suggestive QTLs from mapping various diversity metrics. The table only shows significantly enriched GO terms, according to the Bonferroni- as well as Benjamini-Hochberg (BH) corrected p-values. The study ratio is showing the number of enriched genes within total genes from the respective interval. The population ratio reflects the total number of genes associated with the GO term in the whole annotated genome (total of 59099 genes). The GO enrichment indicates whether the function is overrepresented, ie. enriched (e), or underrepresented, i.e. depleted (p). The enrichment ratio quantifies this effect.

| | QTL (Marker, diversity metric) | GO Term | GO Description | p-val (Bonferroni) | p- val (BH) | GO Enrichment | Study Ratio | Population Ratio | Enrichment Ratio |
|---|---|---|---|---|---|---|---|---|---|
| **Bacteria 2020** | **CHR04_5715673.PC2_wUF** | GO:0036094 | small molecule binding | 1.72E-03 | 4.31E-04 | p | 19/269 | 9539/59099 | 0.0020 |
| | **CHR04_5715673.PC2_wUF** | GO:1901265 | nucleoside phosphate binding | 1.73E-03 | 4.31E-04 | p | 19/269 | 9501/59099 | 0.0020 |
| | **CHR04_5715673.PC2_wUF** | GO:0000166 | nucleotide binding | 1.73E-03 | 4.31E-04 | p | 19/269 | 9501/59099 | 0.0020 |
| | **CHR04_5715673.PC2_wUF** | GO:1901363 | heterocyclic compound binding | 1.73E-03 | 4.31E-04 | p | 19/269 | 9501/59099 | 0.0020 |
| | **CHR13_28627421.PC2_uwUF** | GO:0009628 | response to abiotic stimulus | 1.04E-03 | 5.22E-04 | e | 36/1578 | 595/59099 | 0.0605 |
| | **CHR13_28627421.PC2_uwUF** | GO:0008289 | lipid binding | 1.03E-08 | 1.03E-08 | e | 16/1578 | 73/59099 | 0.2192 |
| | **CHR13_28627421.PC2_uwUF** | GO:0009056 | catabolic process | 1.22E-01 | 4.08E-02 | p | 12/1578 | 1046/59099 | 0.0115 |
| **Bacteria 2022** | **CHR01_7538491.PC2_wUF** | GO:0019725 | cellular homeostasis | 7.68E-03 | 3.84E-03 | e | 13/281 | 732/59099 | 0.0178 |
| | **CHR01_7538491.PC2_wUF** | GO:0042592 | homeostatic process | 7.68E-03 | 3.84E-03 | e | 13/281 | 732/59099 | 0.0178 |
| | **CHR11_569019.PC1_Jaccard** | GO:0038023 | signaling receptor activity | 8.27E-09 | 4.13E-09 | e | 20/505 | 359/59099 | 0.0557 |
| | **CHR11_569019.PC1_Jaccard** | GO:0060089 | molecular transducer activity | 8.27E-09 | 4.13E-09 | e | 20/505 | 359/59099 | 0.0557 |
| | **CHR11_569019.PC1_Jaccard** | GO:0019748 | secondary metabolic process | 1.04E-01 | 3.46E-02 | p | 0/505 | 919/59099 | 0.0000 |
| | **CHR11_569019.PC1_uwUF** | GO:0038023 | signaling receptor activity | 5.07E-05 | 2.54E-05 | e | 24/1186 | 359/59099 | 0.0669 |
| | **CHR11_569019.PC1_uwUF** | GO:0060089 | molecular transducer activity | 5.07E-05 | 2.54E-05 | e | 24/1186 | 359/59099 | 0.0669 |
| | **CHR11_569019.PC1_uwUF** | GO:0005515 | protein binding | 2.25E-02 | 7.51E-03 | e | 101/1186 | 3458/59099 | 0.0292 |
| **Fungi 2022** | **CHR18_18864631. pielou_evenness** | GO:0019748 | secondary metabolic process | 1.55E-02 | 1.55E-02 | e | 53/1953 | 919/59099 | 0.0577 |
